# Chemical modulation of chloroplast de- and re-differentiation reveals a role for the SAL1-PAP retrograde pathway in facilitating plastid transitions

**DOI:** 10.64898/2025.12.11.693750

**Authors:** Pablo Perez-Colao, Jacobo Cruces, Santiago Perez-Rodriguez, Anna Koprivova, Stanislav Kopriva, Aleksandra Skirycz, Jorge Lozano-Juste, Manuel Rodriguez-Concepcion

## Abstract

Plastids are dynamic organelles that remodel their composition, ultrastructure, and function according to developmental and environmental demands. The synthetic molecule X57 induces the conversion of leaf chloroplasts into tocopherol-rich plastids lacking thylakoids and containing proliferating plastoglobules. Removal of X57 triggers chloroplast re-differentiation, enabling precise spatial–temporal dissection of these transitions. X57 directly binds and inhibits the phosphatase SAL1, causing accumulation of its substrate 3′-phosphoadenosine 5′-phosphate (PAP), a retrograde signal that modulates nuclear gene expression. SAL1 inhibition and subsequent PAP accumulation activate a cascade that depletes cytokinins and down-regulates GOLDEN2-LIKE1 (GLK1) and other transcription factors involved in chloroplast biogenesis. SAL1-defective mutants fail to undergo this signaling pathway. The SAL1-PAP-mediated weakening of chloroplast identity preconditions plastids for their eventual conversion into storage-type organelles upon X57-promoted accumulation of tocopherols. After X57 withdrawal, photosynthetic gene expression and chloroplast functions are restored. This framework identifies key molecular mechanisms underlying chloroplast plasticity, a central process in biology.

## INTRODUCTION

Chloroplasts are photosynthetic organelles originally acquired by eukaryotic cells through endosymbiosis with a cyanobacterial ancestor. In plants, they usually differentiate from the proplastids present in meristematic tissues (Jarvis & Lopez-Juez, 2013). The vast majority (over 90%) of the proteins required for plastid assembly and function are nuclear-encoded and must be imported into the organelle after cytosolic synthesis. Therefore, molecular communication between plastids and the nucleus (referred to as retrograde signaling) is crucial for coordinating the transcription of photosynthesis-associated nuclear genes (PhANGs) driving chloroplast differentiation (Chan et al., 2016; Hernández-Verdeja & Strand, 2018). Classic examples of defects in this retrograde communication are found in the *GENOMES UNCOUPLED* (*GUN*) mutants, in which signals from functionally compromised chloroplasts fail to repress PhANG expression, thereby “uncoupling” nuclear transcription from plastid status (Susek et al., 1993; Larkin et al., 2003; Chan et al., 2016). Retrograde signaling, is traditionally divided into: (i) biogenic signals, which operate during chloroplast formation and include intermediates of tetrapyrrole biosynthesis and changes in plastid transcriptional activity; and (ii) operational signals, which respond to environmental or metabolic changes impacting chloroplast function, and involve redox or metabolites cues such as methylerythritol cyclodiphosphate (MEcPP) (Xiao et al., 2012), β-cyclocitral (Ramel et al., 2012) and 3′-phosphoadenosine 5′-phosphate (PAP) (Estavillo et al., 2011). However, this reductionist classification is not mutually exclusive: operational signals can influence chloroplast developmental programs, and biogenic signals can be triggered or modulated by environmental inputs, resulting in considerable overlap between these two categories (Jiang & Dehesh, 2021).

Retrograde signals actively shape the hormonal networks and nuclear transcriptional programs that determine chloroplast activity and fate. Several studies have shown that retrograde pathways modulate responses to hormones such as cytokinins, auxins, gibberellins, and abscisic acid (ABA), thereby influencing both the initiation and maintenance of chloroplast development (Hernández-Verdeja & Strand, 2018; Jiang & Dehesh, 2021; Ortiz-Alcaide et al., 2019). For instance, MEcPP increases salicylic acid accumulation (Xiao et al., 2012), β-cyclocitral modulates jasmonate synthesis (Ramel et al., 2012), and PAP regulates ABA signalling (Phua et al., 2018; Estavillo et al., 2011). This interplay operates as a dynamic feedback loop in which plastid-derived signals influence hormones, which in turn regulate nuclear gene expression to facilitate plastid function, potentially generating new retrograde cues. Thus, hormones influence plastid development both directly and through their interplay with retrograde signaling.

Among them, cytokinins act as positive regulators of chloroplast biogenesis, plastid division and the expression of PhANGs (Cortleven & Schmülling, 2015). Auxins and gibberellins, though classically linked to growth regulation, also modulate chloroplast formation, often acting in opposition or synergy with cytokinins depending on developmental stage and context (Cackett et al., 2022).

At the nuclear level, the integration of retrograde and hormonal inputs converges on transcription factors that operate as master regulators of chloroplast identity. For instance, GOLDEN2-LIKE (GLK) transcription factors, are major drivers of chloroplast developmental programs, with a decisive role in initiating chloroplast biogenesis and maintaining chloroplast function (Hernández-Verdeja & Strand, 2018; Rossini et al., 2001; Waters et al., 2009). GLKs directly activate PhANGs required for thylakoid formation and functional maturation, integrating light-derived signals with information from plastid status and hormonal pathways. Two GLK paralogs are present in *Arabidopsis thaliana*, GLK1 and GLK2. Although they often act redundantly, multiple studies indicate that GLK1 plays the predominant role in many developmental contexts, including response to retrograde and cytokinin signals (Cackett et al., 2022). For instance, *GLK1* overexpression can trigger chloroplast development even in non-photosynthetic tissues, underscoring its central position as a molecular switch for plastid differentiation.

Chloroplasts are dynamic organelles and can de-differentiate to be converted into other plastid types, which often results in their transformation from photosynthetic machines into storage compartments for nutritionally relevant metabolites such as starch (amyloplasts), carotenoids (chromoplasts), fatty acids (elaioplasts) or proteins (proteinoplasts). Furthermore, plastids can interconvert and co-exist with other plastid types in the same cell, illustrating their high degree of plasticity (Choi et al., 2021; Sierra et al., 2023). In some cases, differentiation is reversible. For example, many plant species covert the chloroplasts present in green fruit into carotenoid-rich chromoplasts during ripening. In the process, chloroplasts de-differentiate losing their chlorophyll and thylakoid membranes to accumulate high levels of carotenoid pigments in specialized structures such as lipid droplets (plastoglobules), eventually differentiating into chromoplasts that can re-differentiate back to chloroplasts in some cases (Mesa et al. 2025). In sharp contrast with the wealth of knowledge currently available on chloroplast biogenesis, the molecular mechanisms supporting plastid transitions are less established. This is in part due to the lack of tunable tools to study plastid de- and re-differentiation with spatiotemporal resolution. We previously reported a synthetic compound, X57, that provokes the de-differentiation of chloroplasts into a novel plastid type characterized by the absence of thylakoids and grana and the proliferation of clusters of small lipid bodies or plastoglobules (PG) enriched in tocopherols (Perez-Colao et al., 2025a). Here we show that the effect of X57 is reversible, allowing us to uncover a signaling pathway controlling plastid transitions. Through the use of X57, we found that cytokinins, GLKs and the SAL1-PAP retrograde pathway are fundamental molecular players for chloroplast de-differentiation, weakening chloroplast identity in the first step towards their conversion into tocopherol-overaccumulating plastids.

## RESULTS

### Dynamic and tunable control of chloroplast differentiation by the compound X57

We recently reported the synthetic compound X57 as a new chemical probe to transform chloroplasts into tocopherol-rich plastids (Perez-Colao et al., 2025a). Treatment of *A. thaliana* seedlings with X57 (e.g., by transferring seedlings to X57-supplemented medium) triggers a conspicuous proliferation of clusters of small plastoglobules (PG) and other severe ultrastructural alterations, including the loss of thylakoid membranes, coupled to a reduction in photosynthetic activity **(Fig 1A) (Fig S1)**. Importantly, we found that the effect of X57 is fully reversible. Removal of X57 by transferring seedlings to medium lacking this compound allows a regreening process, characterized by the recovery of normal chlorophyll levels and photosynthetic activity, the gradual reassembly of thylakoid membranes and grana, and a progressive disappearance of PG clusters **(Fig 1A) (Fig S1)**. As a result, only normal leaf chloroplasts were found 6 days after transferring to media without X57 **(Fig 1A)**. Expanding and mature leaves do not contain proplastids but chloroplasts in different developmental stages (Jarvis & Lopez-Juez, 2013). Therefore, we concluded that X57 treatment causes de-differentiation of existing leaf chloroplasts (rather than *de novo* biogenesis of PG-enriched plastids from proplastids) while removal of this synthetic compound results in their re-differentiation back into fully functional chloroplasts.

**Figure 1.**
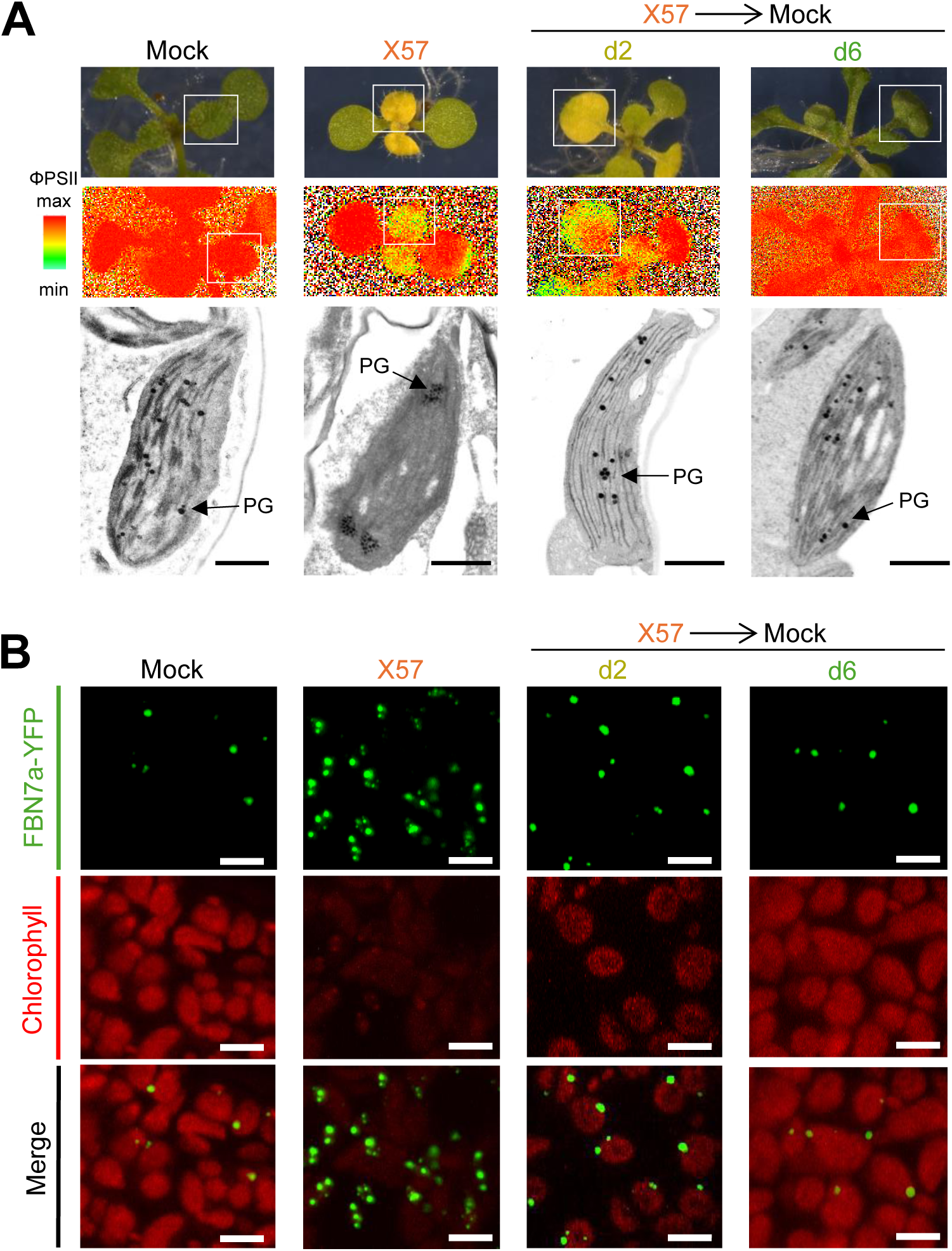
X57 causes chloroplast de-differentiation in a reversible manner. *Arabidopsis thaliana* seedlings were germinated and grown for 5 days on plates containing 0.05% DMSO and then transferred to new plates supplemented with either 0.05% DMSO (mock) or 40 µM X57 for 5 more days. Some X57-treated seedlings were subsequently transferred to mock medium and grown for two (d2) or six (d6) additional days. (A) Representative images of wild-type seedlings and their photosynthetic status (effective quantum yield of photosystem II, ΦPSII). Lower panels correspond to electron microscopy pictures of plastids present in the leaves boxed in the seedling pictures. PG, plastoglobules. Bars, 1 µm. (B) Confocal microscopy analysis of PG distribution in the leaves of transgenic *35S:FBN7a-YFP* seedlings grown as described above. Panels show fluorescence from the PG marker protein FBN7a-YFP (in green) and chlorophyll autofluorescence (in red) in the same microscopy field. Bars, 5 µm.

The dynamic structural changes observed in these experiments were also confirmed *in vivo* using the PG marker line PGL34-YFP (Vidi et al., 2007), herein referred to as *35S:FBN7a-YFP*. Seedlings treated with X57 displayed a strong reduction in chlorophyll autofluorescence together with an increase in FBN7a-YFP signal, in line with the accumulation of PG clusters observed by electron microscopy analysis **(Fig 1)** (Perez-Colao et al., 2025a). Increased chlorophyll autofluorescence and reduced number of FBN7a-YFP fluorescent spots were observed only 2 days after transferring seedlings to medium without X57 **(Fig 1B)**. By day 6 after X57 removal, both chlorophyll autofluorescence and FBN7a-YFP fluorescence were almost identical to those of mock controls never treated with X57 **(Fig 1B).** Altogether, these results open the door to use X57 as a powerful, unprecedented tool to manipulate chloroplast de-and re-differentiation with high precision, hence enabling the analysis of the underlying molecular mechanisms.

### X57 reduces cytokinin levels to promote chloroplast de-differentiation

To look into X57 mode of action, we first analyzed the transcriptomic profile of *A. thaliana* seedlings treated with X57 for 5 h **(Fig 2)**. X57 was previously found to cause changes in the expression of genes for isoprenoid metabolism enzymes and PG-associated structural proteins such as fibrillins (Perez-Colao et al., 2025a). Further KEGG Pathway enrichment analysis of this RNA-seq dataset identified particularly high significance values for zeatin (cytokinin) biosynthesis and plant hormone signal transduction in X57-treated seedlings compared with mock controls **(Fig 2A).** Detailed analysis of genes involved in general cytokinin metabolism **(Fig 2B)** showed that key genes encoding biosynthetic enzymes such as isopentenyltransferases (IPTs) and CYP735As were consistently down-regulated **(Fig 2C)**. By contrast, genes associated with cytokinin degradation and deactivation showed a divergent response. While genes for cytokinin oxidase/dehydrogenases that irreversibly degrade cytokinins (CKXs) were down-regulated, those for UDP-glycosyltransferases that conjugate cytokinins to glucose to inactivate them (UGTs) were up-regulated **(Fig 2C).**

**Figure 2.**
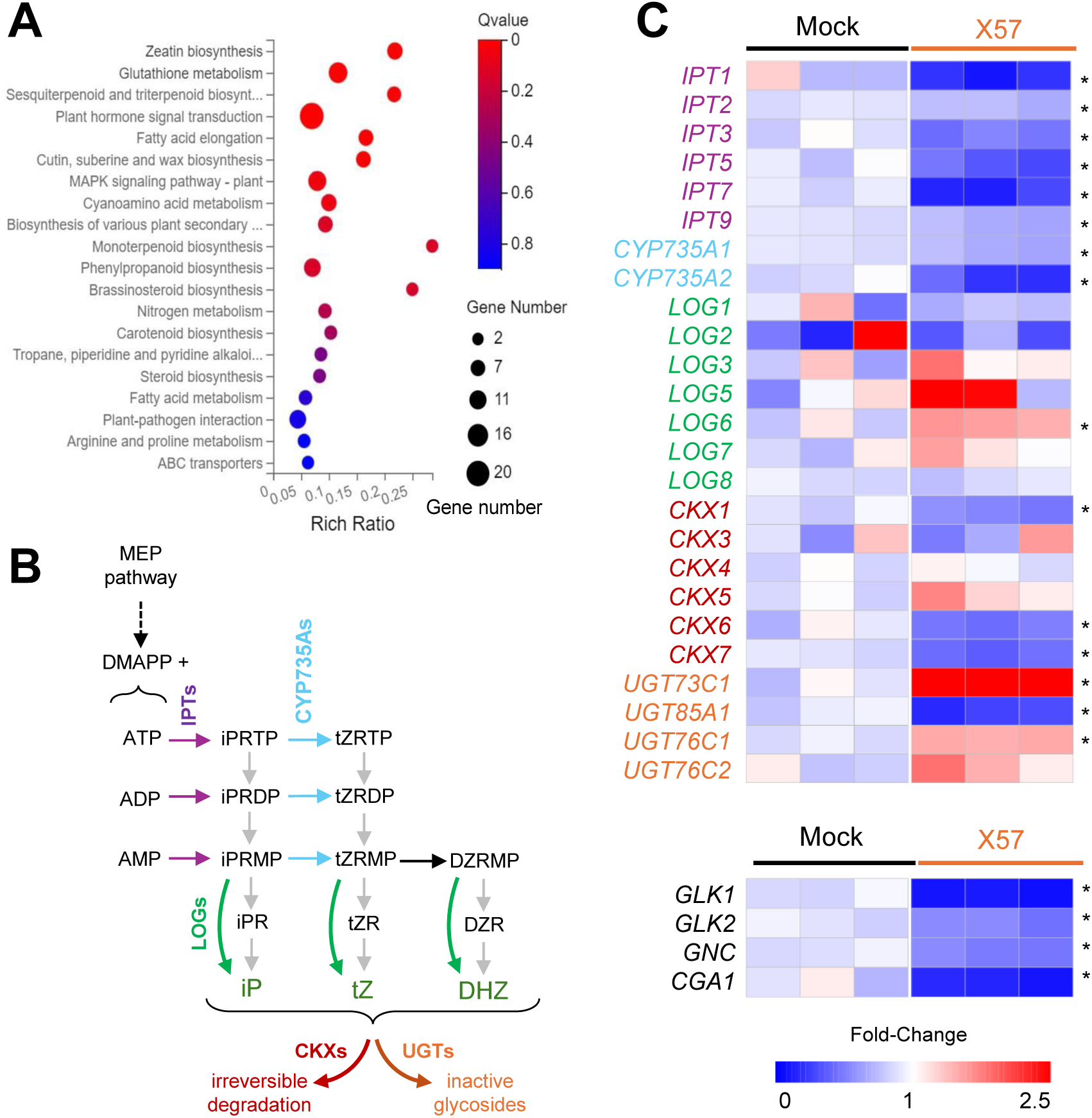
X57 represses expression of genes involved in cytokinin biosynthesis and chloroplast biogenesis. *A. thaliana* seedlings were germinated and grown for 5 days on plates containing 0.05% DMSO and then transferred to liquid medium supplemented with either 0.05% DMSO (mock) or 100 µM X57. Samples for RNA-seq were collected 5 h after transfer. (A) KEGG pathway enrichment analysis of differentially expressed genes (DEGs) in X57-treated versus mock-treated seedlings. Qvalue represents the significance value of enrichment (values lower than 0.05 correspond to significant enrichment). The size of the bubbles represents the number of DEGs annotated to a particular term. (B) Schematic diagram of cytokinin metabolism. Enzymes are indicated with colors: IPTs, isopentenyl transferases; CYP735As, cytochrome P450 family 735A; LOGs, LONELY GUY phosphoribohydrolases; CKXs, cytokinin oxidases / dehydrogenases; UGTs, UDP-glycosyltransferases. Active cytokinins and marked in green: iP isopentenyladenine; tZ, trans-zeatin; DHZ, dihydrozeatin. (C) Heatmaps representing transcript abundance of genes involved in cytokinin biosynthesis (IPTs, CYP735As, LOGs), degradation (CKXs), and inactivation (UGTs), as well as genes encoding transcription factors regulating chloroplast biogenesis (lower panel). Fold-Change represents normalized values using the row_scale method of the pheatmap R package, and columns correspond to biological replicates. Asterisks indicate significant changes in X57 compared to mock samples (DESeq2; FDR ≤ 0.05). Gene accessions and TPM values are listed in Table S2.

To confirm whether the observed down-regulation of biosynthetic genes together with the up-regulation of genes for cytokinin deactivation resulted in lower levels of biologically active hormones, we quantified the contents of the three main active cytokinins in *A. thaliana* seedlings treated with X57 for 5 h **(Fig 3A)**. Hormone profiling revealed a sharp reduction in isopentenyladenine (iP) and, to a lower extent, dihydrozeatin (DHZ), whereas the decrease in trans-zeatin (tZ) was not statistically significant at this stage **(Fig 3A)**. Importantly, no major changes were detected for other phytohormones in the same samples **(Fig S2)**. Measurement of cytokinin levels in seedlings exposed to X57 for 48 h, when the phenotypic effects of X57 became clearly visible **(Fig S3)**, showed that all three active cytokinins (iP, DHZ, and tZ) were strongly reduced in X57-treated plants compared with mock controls **(Fig 3A)**. These results indicate that X57 causes a rapid reprogramming of cytokinin metabolism at the level of gene expression that causes a depletion of active hormones.

**Figure 3.**
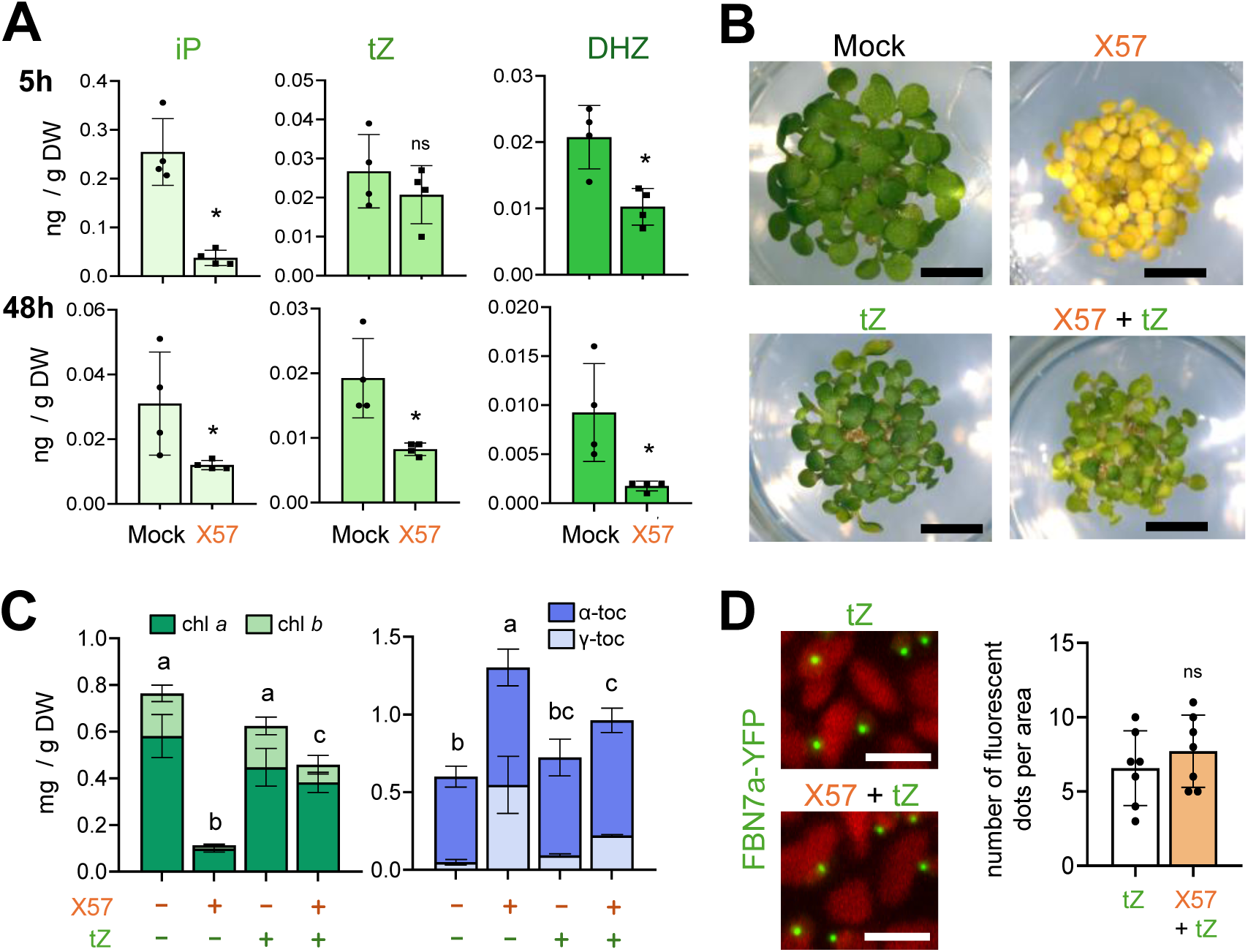
Cytokinin depletion is required for X57-mediated chloroplast de-differentiation. (A) Levels of active cytokinins in *A. thaliana* seedlings germinated and grown for 5 days on plates containing 0.05% DMSO and then transferred to liquid medium supplemented with either 0.05% DMSO (mock) or 100 µM X57 for 5 and 48 h. Mean and SD of four independent samples (n=4) are represented. Statistically significant differences relative to mock samples (t-test, *p* < 0.05) are represented with asterisks (ns, not significant). DW, dry weight. (B) Representative images of seedlings germinated and grown for 5 days in solid medium supplemented with 0.05% DMSO (mock), 25 µM X57, 5 µM trans-zeatin (tZ), or both 25 µM X57 and 5 µM tZ. Bars, 1 mm. (C) Levels of chlorophylls (*a* and *b*) and tocopherols (α and γ) in seedlings treated (+) or not (-) with the indicated compounds as described in (B). Mean and SD of three independent samples (n=3) are represented. Letters indicate statistical significance (*p* <0.05) according to one-way ANOVA followed by Tukey’s post hoc test of total chlorophyll or tocopherol contents. (D) Confocal microscopy analysis of PG distribution in transgenic *35S:FBN7a-YFP* seedlings grown on media supplemented with tZ or both X57 and tZ, as described in (B). Panels show fluorescence from the PG marker protein FBN7a-YFP (in green) and chlorophyll autofluorescence (in red). Bars, 5 μm. Plot shows the quantification of PGs (green fluorescent dots) in microscopy images. Mean and SD of seven different leaf areas (n=7) are shown. No statistically significant differences (t-test, *p* < 0.05) were found (ns, not significant).

Next, we tested whether cytokinin depletion was required for the X57-triggered chloroplast de-differentiation phenotype by growing seedlings in the presence of exogenously supplied tZ. Addition of this cytokinin partially complemented the golden phenotype produced by X57 **(Fig 3B)** by restoring chlorophyll and tocopherol contents to levels close to those of the mock condition or cytokinin treatment alone **(Fig 3C)**. Analysis of PG proliferation with the *FBN7a-YFP* marker line further confirmed that tZ supplementation prevented X57-triggered chloroplast de-differentiation **(Fig 3D)**.

### GLK transcription factors orchestrate chloroplast de- and re-differentiation

Since cytokinins exert their function largely through transcriptional regulation of chloroplast developmental programs, we next examined whether genes encoding transcription factors downstream of cytokinin signaling (Cackett et al., 2022; Chan et al., 2016) were misregulated in X57-treated seedlings. Strikingly, X57 treatment caused a coordinated repression of the four main regulators of chloroplast biogenesis: *GLK1* (AT2G20570)*, GLK2* (AT5G44190)*, GATA NITRATE-INDUCIBLE CARBON-METABOLISM-INVOLVED* (*GNC*; AT5G56860), and *CYTOKININ-RESPONSIVE GATA FACTOR 1* (*CGA1*; AT4G26150) **(Fig 2C).** *GLK1*, which has a central role in chloroplast biogenesis, was selected to test the functional relevance of its repression by X57. We hypothesized that if *GLK1* down-regulation is required for the X57-triggered loss of chloroplast identity, constitutive overexpressing GLK1 (i.e., in a manner independent of X57 and cytokinins) could counteract the chloroplast de-differentiation phenotype induced by X57. To test this hypothesis, we initially used transgenic *A. thaliana* lines carrying a *35S:GLK1* construct, but realized that they were not true overexpressors because the expression of known GLK1 target genes such as *LHCB1B1/LHCB1.4* (AT2G34430), *CAB3/ LHCB1.2* (AT1G29910), and *PETE1* (AT1G76100) was not induced. We therefore decided to use transient expression in *Nicotiana benthamiana* leaves (to ensure high expression levels). Leaves co-agroinfiltrated with constructs *35S:GLK1* or/and *35S:FBN7a-GFP* (to monitor PG proliferation) were subsequently treated with either X57 or a mock solution. While infiltration with X57 alone reduced chlorophyll accumulation, increased tocopherol levels, and promoted PG proliferation, co-infiltration with GLK1 abolished all these X57-dependent phenotypes **(Fig 4A-B)**, confirming that high GLK1 levels prevent X57-mediated chloroplast de-differentiation.

**Figure 4.**
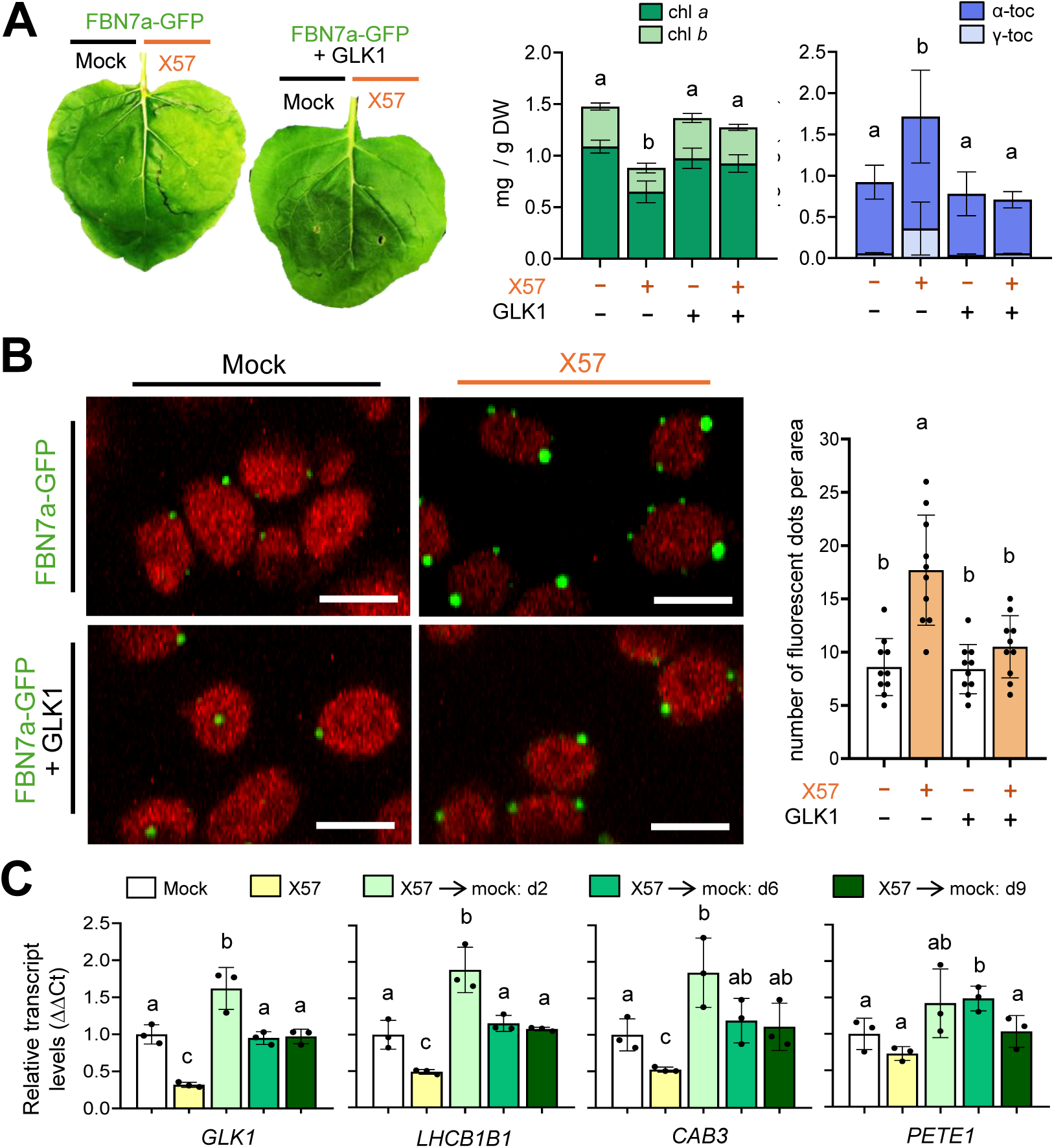
Chloroplast de-differentiation requires down-regulation of GLKs. (A) *Nicotiana benthamiana* leaves were agroinfiltrated with constructs to produce the PG marker FBN7a–GFP alone (control) or together with the *A. thaliana* transcription factor GLK1, and 24 h later they were treated with 0.1% DMSO (mock) or 100 µM X57. Representative leaves harvested 4 days after agroinfiltration (3 days after treatment) are shown. Plots represent levels of chlorophylls (*a* and *b*) and tocopherols (α and γ) in these leaves. Mean and SD of three independent samples (n=3) are represented. DW, dry weight. (B) Confocal microscopy analysis of PG distribution in *N. benthamiana* leaves treated as described in (A) and collected 3 days after agroinfiltration (2 days after treatment). Panels show FBN7a-GFP fluorescence (in green) and chlorophyll autofluorescence (in red). Bars, 5 μm. Plot shows the quantification of PGs (green fluorescent dots) in microscopy images. Mean and SD of ten different leaf areas (n=10) are shown. (C) *A. thaliana* seedlings germinated and grown for 5 days on mock plates were transferred to new plates supplemented with either 0.05% DMSO (mock) or 40 µM X57 for 5 more days, and then some X57-treated seedlings were transferred back to mock plates and grown for two (d2), six (d6), or nine (d9) additional days. Plots show RT-qPCR analysis of transcript levels for the indicated genes in the samples, relative to mock controls. Mean and SD of three independent replicates (n=3) are shown. In all plots, letters indicate statistically significant differences (*p*<0.05) according to two-way ANOVA followed by Tukey’s post hoc test.

Once we found that *GLK1* down-regulation was required for chloroplast de-differentiation, we hypothesized that restoration of *GLK1* expression might be required for chloroplast re-differentiation during the regreening process following X57 removal. As a first estimate, we quantified transcript levels of *GLK1* and known target genes in X57-treated *A. thaliana* seedlings before and after transferring them to X57-depleted medium on day 0. RT-qPCR analysis revealed that *GLK1* and its target genes *LHCB1B1*, *CAB3*, and *PETE1*, which encode light-harvesting chlorophyll a/b-binding proteins and components of the photosynthetic electron transport chain, were strongly repressed in X57-grown seedlings at day 0 **(Fig 4C)**. However, upon transfer to mock medium, *GLK1* and its downstream PhANGs showed a peak induction at day 2 and returned to basal levels by day 9 **(Fig 4C)**. Altogether, our data confirm that GLK transcription factors promote re-differentiation (a role well established in the literature), but they also prevent chloroplast de-differentiation.

### X57 targets SAL1 to enhance PAP accumulation

To determine the molecular mechanism by which X57 triggers chloroplast de-differentiation, we searched for direct molecular targets using isothermal shift assay (iTSA), a mass spectrometry–based approach that detects ligand-induced changes in protein thermal stability allowing for proteome-wide identification of small molecule targets (Ball et al., 2020; Schlossarek et al., 2022). To minimize false positives and increase the likelihood of identifying a direct protein target bound by X57, we synthesized a structurally related dead analog. The resulting analog, named X56, shared the core structure of X57 but it did not trigger changes in chlorophyll or tocopherol levels and it also failed to induce PG proliferation **(Fig. S4)**.

Protein extracts from *A. thaliana* seedlings were incubated with X57 or with the dead analog X56 as a negative control and subjected iTSA. Proteins differentially accumulated in the soluble fraction of the samples spiked with X57 vs X56 (putative X57 targets) were identified by mass spectrometry. Among the 3092 proteins detected in the iTSA assay, 179 showed a differential abundance between X57 and X56 treatments **(Table S1)**. From this subset, we filtered for candidates with known functions in chloroplast development or/and hormone signaling. Among them, we identified the protein SAL1 (AT5G63980), a 3′(2′),5′-bisphosphatase that dephosphorylates 3′-phosphoadenosine 5′-phosphate (PAP) to adenosine monophosphate (AMP) and inorganic phosphate. Inactivation of SAL1 results in the accumulation of PAP, a retrograde signal that translocates to the nucleus, where it inhibits RNA degrading exoribonucleases to modulate the expression of genes involved in a variety of plant responses, including those related to hormones (Chan et al., 2016; Estavillo et al., 2011; Phua et al., 2018). We next confirmed the binding of X57 to SAL1 *in vitro* by nano differential scanning fluorimetry (nanoDSF) (Gao et al., 2020; Magnusson et al., 2018). While the inactive analogue X56 had no effect on SAL1 unfolding (Tm 61.2°C in X56 vs. 61.1°C in mock-treated controls), incubation with X57 consistently produced a shift on the protein Tm to 59.5°C, supporting a direct interaction between SAL1 and X57 **(Fig 5A)**. To further validate this interaction and determine the binding affinity of X57 for SAL1, we performed microscale thermophoresis (MST) assays (Magnusson et al., 2018; Seidel et al., 2013) using fluorescently labelled SAL1. The results confirmed direct binding of X57 to SAL1 with a dissociation constant K_d_ = 2.07 µM, while no binding was observed for X56 **(Fig 5B)**. Together, our results provide strong evidence supporting the specific binding of X57 to SAL1 *in vitro*.

**Figure 5.**
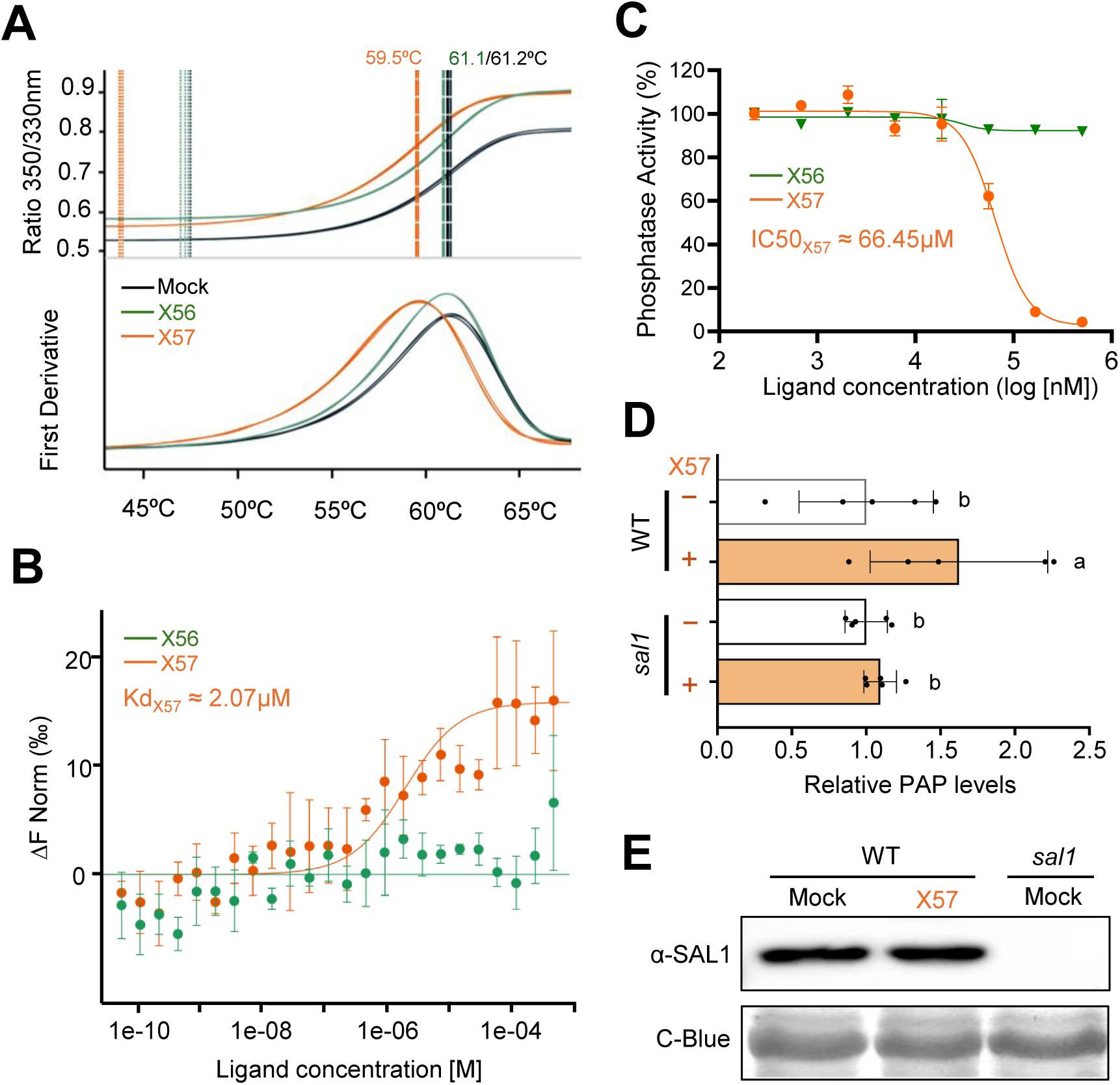
Direct binding of X57 to SAL1 represses its activity and results in enhanced PAP accumulation. (A) Thermal stability of purified His-tagged SAL1 assessed by nano differential scanning fluorimetry (nanoDSF). Protein samples (2 µM) were incubated with 0.1% DMSO (mock), 100 µM X57, or 100 µM X56 (dead analog), and subjected to a linear temperature ramp. Intrinsic fluorescence at 330 and 350 nm was recorded and plotted as raw unfolding curves and first derivative plots. (B) Microscale thermophoresis (MST) analysis of fluorescently labeled His-tagged SAL1 mixed with serial dilutions of ligand (X57 or X56). (C) Phosphatase assay of His-tagged SAL1 using para-methylumbelliferyl phosphate (pMUP) as a fluorogenic substrate. Purified protein (50 nM) was pre-incubated with X57, X56, or a mock soluction for 20 min prior to substrate addition. (D) PAP contents in *A. thaliana* wild-type (WT) and mutant *sal1* seedlings germinated and grown for 5 days in medium supplemented with 0.05% DMSO (-) or 25 µM X57 (+). Levels are represented relative to those in mock samples of the same genotype. See Figure S5 for absolute values. (E) Immunoblot analysis of SAL1 protein levels in the samples used in (D). Coomassie Brilliant Blue (C-Blue) staining is shown as a loading control. Plots represent mean and SD of n=3 (A–C) or n=5 (D) independent biological replicates. Letters in (D) indicate statistically significant differences (*p*<0.05) according to one-way ANOVA followed by Tukey’s post hoc test.

To test whether the interaction with X57 modulates the activity of SAL1, we quantified the phosphatase activity of the enzyme *in vitro* using the fluorogenic substrate pMUP (para-methylumbelliferyl phosphate). The results conclusively showed that X57 was able to inhibit SAL1 phosphatase activity *in vitro*, whereas the inactive analogue X56 had no significant effect at the same concentrations **(Fig. 5C)**. The observed X57-dependent inhibition followed a dose-dependent profile, showing an IC₅₀ of 66.45 µM **(Fig. 5C)**. Whether X57 was also able to inhibit SAL1 activity *in vivo* was estimated by measuring PAP levels in *A. thaliana* seedlings treated or not with X57. In agreement with the inhibitory effect of X57 on SAL1 activity detected *in vitro*, a significant increase in PAP levels without changes in SAL1 protein accumulation was observed in X57-treated wild-type (WT) seedlings **(Fig. 5D-E) (Fig. S5)**. In SAL1-defective mutants, PAP levels were much higher but they remained unchanged in the presence of X57 **(Fig. 5D) (Fig. S5)**, confirming that altered PAP accumulation in WT seedlings exposed to X57 resulted from reduced SAL1 function. Furthermore, we tested whether increased PAP levels resulting from X57 application could activate retrograde signaling by monitoring the expression of genes known to be regulated by the SAL1-PAP pathway (Khin et al., 2025) such as *ANAC013* (AT1G32870), *AOX1a* (AT3G22370), *APX2* (AT3G09640), *CPK32* (AT3G57530), *RCD1* (AT1G32230), and *SOT12* (AT2G03760). Some of these genes were indeed up-regulated in X57-treated WT seedlings **(Fig. S6)**. These results together demonstrate that direct binding of X57 to SAL1 causes a reduction in its catalytic activity, resulting in increased accumulation of the retrograde signal molecule PAP and subsequent rewiring of nuclear gene expression.

### SAL1 is required for X57-triggered chloroplast de-differentiation

If X57 inhibition of SAL1 activity is physiologically relevant for the control of chloroplast de-differentiation, it would be expected that *sal1* seedlings remained greener than WT controls when treated with X57. Indeed, SAL1-defective seedlings germinated and grown in the presence of X57 did not acquire the characteristic golden phenotype and remained green even at concentrations that completely bleached the WT controls **(Fig 6A**). Consistently, chlorophyll drop in X57-germinated WT seedlings was attenuated in *sal1* seedlings, whereas the X57-mediated up-regulation of tocopherol accumulation was similar in WT and *sal1* samples **(Fig. 6A)**. A similar greening effect was observed in seedlings germinated without X57 and then transferred to media supplemented with different concentrations of X57, as the percentage of golden leaves generated after the transfer remained lower in *sal1* seedlings compared to WT controls **(Fig. 6B)**. At the ultrastructural level, the chloroplasts of WT and *sal1* leaves were similar under mock conditions **(Fig. 6C)**. In leaves from seedlings transferred to X57, however, *sal1* chloroplasts retained thylakoids and few PG in strike contrast with the thylakoid-devoid and PG-rich plastids found in the WT **(Fig. 6C)**.

**Figure 6.**
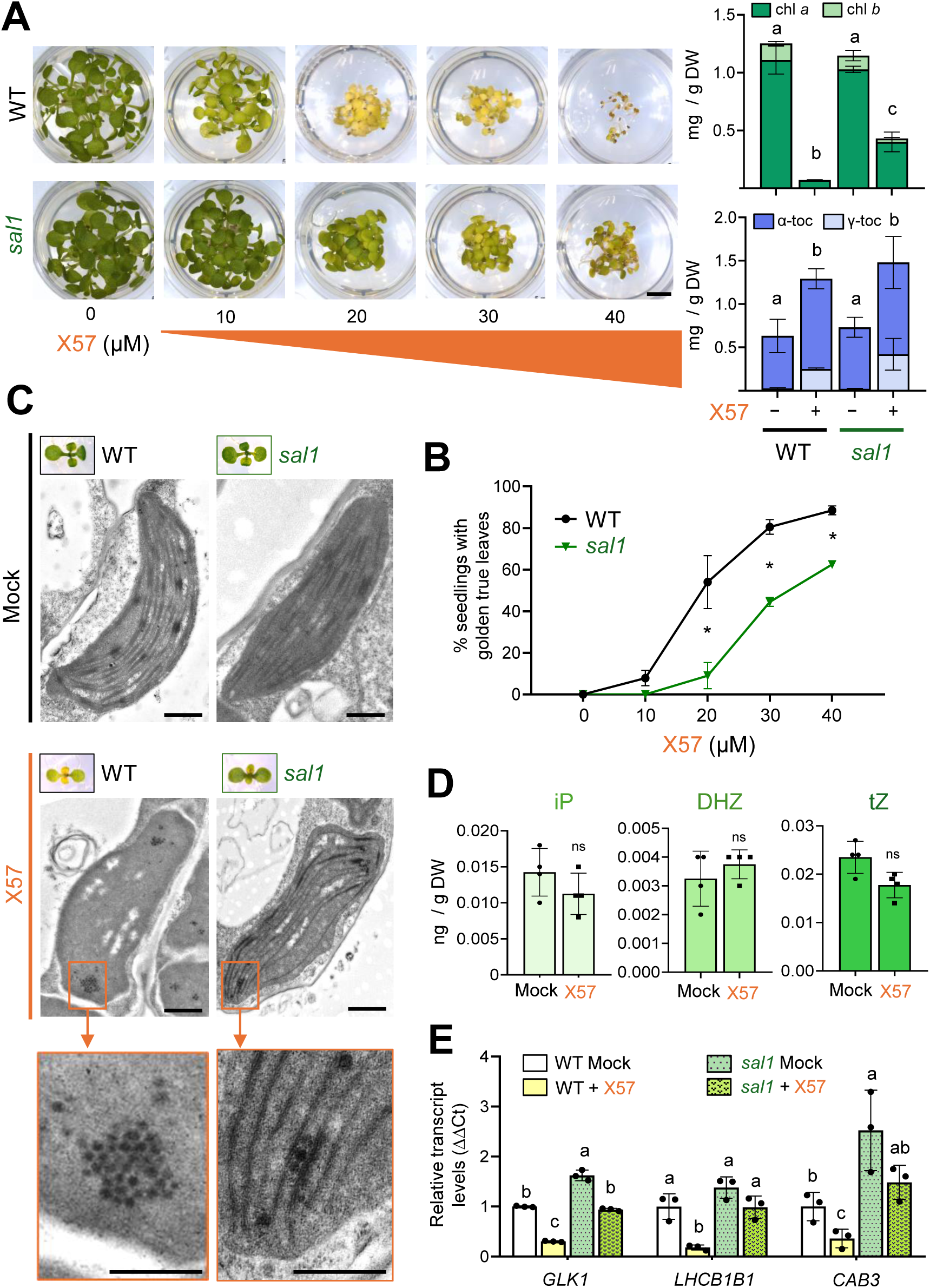
X57-mediated chloroplasts de-differentiation requires regulation of SAL1 activity. (A) Representative images of *A. thaliana* wild-type (WT) and *sal1* seedlings germinated and grown for 7 days on medium supplemented with the indicated concentrations of X57. Plots represent levels of chlorophylls (*a* and *b*) and tocopherols (α and γ) in seedlings from 0 µM (-) and 25 µM X57 (+) treatments. Mean and SD of three independent samples (n=3) are represented. DW, dry weight. (B) WT and *sal1* seedlings germinated and grown for 5 days in mock medium were transferred to fresh medium supplemented with the indicated concentrations of X57. After 5 days, the proportion of seedlings with chlorophyll-depleted (i.e., golden) leaves was monitored. Mean and SD of three biological replicates (n=3) are represented. (C) Representative plastids from the true leaves of the seedlings shown in the pictures (bars, 1 µm). Lower panels correspond to magnifications of the boxed areas of plastids from X57-treated seedlings (bars, 0.5 µm). (D) Levels of active cytokinins in *sal1* seedlings germinated and grown for 5 days on plates containing 0.05% DMSO and then transferred to liquid medium supplemented with either 0.05% DMSO (mock) or 100 µM X57 for 5 h. Mean and SD of four independent samples (n=4) are represented. (E) RT-qPCR analysis of transcript levels for the indicated genes in WT and *sal1* seedlings germinated and grown in medium supplemented with 0.05% DMSO (mock) or 25 µM X57 for 7 days, relative to WT mock controls. Mean and SD of three independent replicates (n=3) are shown. Statistically significant differences are represented with letters (one-way ANOVA followed by Tukey’s post hoc test, *p*<0.05) or asterisks (t-test: *, *p*<0.05; ns, not significant).

Next, we tested whether SAL1 was required to trigger the molecular mechanism by which X57 promotes chloroplast de-differentiation, i.e., the down-regulation of cytokinins, GLK1, and PhANGs. Levels of active cytokinins dropped soon (5h) after X57 treatment in WT seedlings **(Fig. 3A)** but remained unchanged in *sal1* seedlings **(Fig. 6D)**. The X57-mediated decrease in the expression of *GLK1* and target PhANGs such as *LHCB1B1* and *CAB3*, was also attenuated in the mutant **(Fig 6E)**. We therefore concluded that SAL1 is required for efficient chloroplast de-differentiation induced by X57 via cytokinins and GLK-mediated gene expression. To investigate whether other retrograde signaling pathways besides the SAL1–PAP pathway could also mediate the described effects of X57, we analyzed the phenotype of different mutants defective in plastid gene expression (*gun1* and *rif10*), tetrapyrrole metabolism (*gun2, gun3, gun4,* and *gun5*), and MEcPP synthesis (*csb3*). None of these mutants displayed altered sensitivity to X57 **(Fig. S7)**, highlighting the specificity of the SAL1-PAP pathway on the X57-mediated regulation of chloroplast de-differentiation.

## DISCUSSION

Chloroplasts are central to photosynthesis, fixing atmospheric CO₂ and producing O₂. The establishment, maintenance, and remodeling of chloroplast identity depend on tight coordination between nuclear transcriptional programs and plastid-derived signals. Nuclear–plastid communication is also required for chloroplast conversion into other plastid types specialized in the accumulation of metabolites such as starch (amyloplasts), fatty acids (elaioplasts), proteins (proteinoplasts) or carotenoid pigments (chromoplasts). Yet, dissecting the signaling mechanisms that govern chloroplast plasticity has remained difficult because genetic perturbations lack temporal reversibility. Here we introduce X57, a small-molecule probe that enables the precise manipulation of chloroplast de- and re-differentiation. Treatment with X57 transiently converts functional chloroplasts into PG-enriched plastids that lack thylakoid membranes, whereas its removal restores chloroplast structure and photosynthetic capacity completely **(Fig. 1) (Fig. S1)**. This reversibility distinguishes X57 from genetic mutants with irreversible chloroplast defects (Myouga et al., 2013) and from chemicals inducing permanent damage such as norflurazon, fosmidomycin, or lincomycin (Ortiz-Alcaide et al., 2019; Sauret-Güeto et al., 2006; Susek et al., 1993). In summary, X57 stands as a unique chemical tool that has allowed us to unveil key molecular players driving chloroplast transitions.

Thermoproteomic, biochemical, and genetic analyses identified SAL1 as a molecular target of X57. Binding of X57 to SAL1 inhibits its phosphatase activity **(Fig. 5C)**, elevates the retrograde signaling molecule PAP *in vivo* **(Fig. 5D)**, and rapidly triggers changes in nuclear gene expression **(Fig. S6)**. The molecular, biochemical, and phenotypic effects of X57 are strongly attenuated in *sal1* seedlings **(Fig. 6)** but not in mutants defective in other retrograde pathways **(Fig. S7)**, demonstrating that the SAL1–PAP signaling pathway mediates X57-induced chloroplast de-differentiation. Our results reveal a functional connection between SAL1–PAP signaling and cytokinin metabolism in agreement with transcriptomic alterations on cytokinin signaling reported for *sal1* mutants (Chan et al., 2016; Estavillo et al., 2011). X57 rapidly depletes active cytokinins in WT seedlings **(Fig.3A)** but not in the *sal1* mutant **(Fig. 6D)**. Furthermore, exogenous cytokinin supplementation restores chlorophyll content and suppresses PG proliferation in X57-treated WT seedlings **(Fig. 3)**, establishing cytokinin depletion as a causal trigger of chloroplast de-differentiation. Cytokinins promote the differentiation of chloroplasts by regulating the expression of both nuclear and plastid genes (Cortleven & Schmülling, 2015; Chan et al., 2016; Cackett et al., 2022). Among them, cytokinins induce expression of genes for key nuclear transcription factors involved in chloroplast biogenesis such as GLK1, GLK2, GNC, and CGA1, all of which were strongly repressed by X57 **(Fig. 2)**. This repression is required for X57-mediated chloroplast de-differentiation **(Fig. 4)**, suggesting that GLK transcription factors counteract the de-differentiation program imposed by X57. The induction of GLK1 and its target PhANGs when chloroplast re-differentiate following X57 removal **(Fig. 4C)** and further underscores their role in restoring chloroplast identity. Together, our results reveal a reversible SAL1–PAP/cytokinin/GLK axis that modulates chloroplast identity **(Fig. S8)**. SAL1 inhibition (e.g., by treating with X57) elevates PAP, represses cytokinin biosynthesis, down-regulates GLK activity, and reduces *PhANG* expression, hence weakening chloroplast identity. Upon restoration of SAL1 activity (e.g., by removing X57), *GLK1* and *PhANG* expression recover, allowing chloroplast to regain normal photosynthetic functions.

The SAL1-PAP pathway, conserved in all land plants and their algal ancestors (Zhao et al., 2019), mediates operational retrograde signaling in response to redox stress cues that challenge normal chloroplast functioning (Chan et al., 2016; Jiang & Dehesh, 2021). When redox balance is disrupted, SAL1 activity decreases and PAP accumulates, eventually moving to the nucleus to modulate gene expression by inhibiting RNA-degrading nuclear exoribonucleases (XRNs) and altering post-transcriptional gene silencing, mRNA turnover, and transcription (Estavillo et al., 2011). It remains to be determined whether SAL1 activity is dynamically regulated during natural plastid transitions, but our results suggest that challenging chloroplast identity is the first key requirement for chloroplasts to transition into a different plastid type. A second essential requisite appears to be the accumulation of metabolites (e.g., starch, fatty acids, proteins or carotenoids) promoting the transformation of photosynthetic membranes into storage structures (Llorente et al., 2020; Li et al., 2025). In the case of X57, the SAL1-mediated down-regulation of cytokinin levels and eventually GLK activity and *PhANG* expression makes chloroplasts competent for de-differentiation, which takes place thanks to the SAL1-independent up-regulation of tocopherol accumulation **(Fig. 6A)** and eventual PG development (Perez-Colao et al., 2025a). Without sustained tocopherol increases, chloroplasts remain stressed but do not transform, explaining the absence of the X57-dependent golden phenotype in untreated *sal1* mutant seedlings or WT seedlings treated with other inhibitors of SAL1 (Khin et al., 2025). Conversely, tocopherol overaccumulation alone is insufficient to cause chloroplast de-differentiation (Li et al., 2010; Lu et al., 2013).

Our results suggest a two-step paradigm across chloroplast conversions: (1) chloroplast conditioning *via* reduction or loss of photosynthetic capacity, followed by (2) ultrastructural rearrangements driven by enhanced levels of specific metabolites. How metabolic perturbations differentially shape nuclear programs to eventually support ultrastructural changes associated to particular plastid types remains unknown. Addressing this important challenge would require new tools able to uncouple the different signaling pathways involved. In this context, X57 provides a tunable chemical tool for the temporal dissection of chloroplast de-differentiation but also re-differentiation. Besides offering novel experimental evidence that chloroplast differentiation represents a dynamically maintained equilibrium, rather than a terminally fixed developmental commitment, the use of X57 has allowed to define a new molecular SAL1-PAP/cytokinin/GLK axis that offers a framework for manipulating chloroplast plasticity to eventually obtain more nutritive and stress-resilient crops. Because PAP also influences stress responses and hormone networks, the ability of X57 to control its levels chemically opens new avenues for dissecting the multifaceted roles of retrograde communication in plant adaptation.

## MATERIAL AND METHODS

### Plant material, chemical treatments, and growth conditions

All *Arabidopsis thaliana* lines used in this study were in the Columbia (Col-0) background, including the PGL34-YFP marker line, here referred to as *35S:FBN7a-YFP* (Vidi et al., 2007); *csb3* (Gil et al., 2005); *gun1, gun2* and *gun3* (Susek et al., 1993); *gun4* (Larkin et al., 2003)*; gun5* (Mochizuki et al., 2001); and *rif10* (Sauret-Güeto et al., 2006). Two *sal1* mutant alleles were used: *ron1-1* (Robles et al., 2009) for most experiments **(Fig. 6) (Fig. S5)** and the *fry1* line SALK_020882 (Ashykhmina et al., 2019) for the comparison with other retrograde mutants **(Fig. S7)**. Seeds were surface-sterilized with 70% ethanol for 5 min, followed by 100% ethanol for 1 min, and then air-dried. Unless indicated otherwise, sterile seeds were directly sown onto solid Murashige and Skoog (MS) medium with 1% sucrose in 24-well square cell culture plates or 9 cm Petri dishes. When indicated, the medium was supplemented with different concentrations of X56 or X57, or with 0.05% DMSO as a mock control. X57 and X56 **(Fig. S4)** belong to a proprietary screening library from GalChimia and were provided directly by them and diluted in DMSO to prepare a 50 mM stock solution. Trans-zeatin riboside (Sigma-Aldrich) was diluted in water and added to the medium in some experiments. Stratification was carried out at 4 °C in darkness for 3 days. Then, plates were incubated in a growth chamber at 22 °C under long-day (LD) conditions of 8 h of darkness and 16 h of fluorescent white light, with a Photosynthetic Photon Flux Density (PPFD) of 100 µmol m⁻² s⁻¹. For transfer experiments, seedlings were germinated on mock MS medium and grown for 5 to 7 days under LD conditions. Then, they were transferred to fresh X57-supplemented or mock plates and incubated under LD for several days. For regreening assays, X57-transferred seedlings were moved back to mock medium and samples were collected the day of transfer or several days afterwards. *Nicotiana benthamiana* plants used for transient expression assays were grown in a greenhouse under a LD photoperiod of 8h of darkness (at 21 °C) and 16 h of light (at 26 °C) with a PPFD of 100 µmol m⁻² s⁻¹.

### Photosynthetic measurements

Effective quantum yield of PSII (ΦPSII) was measured and imaged using a Handy-GFP fluorometer (Photon Systems Instrument) on intact true leaves from *A. thaliana* seedlings growing on plates. Prior to each measurement, seedlings were dark-adapted for 15 min to ensure full relaxation of PSII reaction centers. After the dark period, minimal fluorescence (F₀) was recorded using a weak modulated measuring beam, following the instrument protocol parameters (FoDuration = 5 s; FoMeasure = 1 s). A saturating pulse of approximately 800 ms was then applied to determine maximal fluorescence (Fₘ) in dark-adapted state. Immediately afterwards, leaves were exposed to actinic light of 10 μmol photons m⁻² s⁻¹, and steady-state fluorescence (Fₛ) was recorded once fluorescence stabilization was reached. Additional saturating pulses of 800 ms were delivered under actinic illumination to obtain maximal fluorescence in the light-adapted state (Fₘ′). ΦPSII was calculated as (Fₘ′ – Fₛ)/Fₘ′. For each experimental condition, at least five biological replicates (seedlings from different plates) were analyzed.

### Metabolite extraction and quantification

Chlorophylls and tocopherols were extracted and quantified as described (Perez-Colao et al., 2025a). PAP was determined as described (Ashykhmina et al., 2019). In brief, lyophilized plant material was homogenized with 500 μl of hot (80°C) extraction buffer (62 mM citric acid, 76 mM K_2_HPO_4_, pH 4) and after incubation for 5 min at 80°C and cooling on ice for 15 min the samples were centrifuged. Supernatants were derivatized with chloracetaldehyde for 40 min at 60°C. After cooling, centrifugation, and dilution 1:1 with water, the samples were analyzed using reversed-phase HPLC on Hyperclone C18 BDS column and a gradient of acetonitrile in tetrabutylammoniumbisulfate buffer (5.7 mmol TBAS and 30.5 mmol K_2_HPO_4_ pH 5.8). The derivatized nucleotides were detected by a Dionex RF 2000 fluorescence detector and quantified using external standards.

### Hormone measurements

Plant growth regulators including acidic hormones (gibberellins, indoleacetic acid, jasmonic acid, abscisic acid, and salicylic acid) and basic hormones (cytokinins, including dihydrozeatin, isopentenyladenine, and trans-zeatin), were identified and quantified at the IBMCP’s Hormone Quantification Platform essentially as described (García et al., 2020). Briefly, 50–100 mg of freeze-dried tissue was suspended in 80% methanol containing 1% acetic acid, stirred for 1 h at 4°C, and centrifuged at 14,000 rpm for 4 min at 4°C. The supernatant was stored overnight at −20°C and centrifuged again (14,000 g, 4 min, 4°C). The clarified extract was dried in a vacuum concentrator and resuspended in 1% acetic acid. Samples were consecutively passed through an Oasis HLB reverse-phase column (30 mg; Waters) and an Oasis MCX cation exchanger. Acidic hormones were eluted with methanol, and basic hormones with 60% methanol containing 5% aqueous ammonia. The eluates were dried and reconstituted in 5% acetonitrile, 1% methanol, and 1% acetic acid. Separation was performed by ultra-high-performance liquid chromatography (UHPLC) on an Accucore C18 column (2.6 µm, 2.1 × 100 mm; Thermo Scientific) using a linear gradient of 2–55% acetonitrile containing 0.05% acetic acid over 21 min at 0.4 mL min⁻¹. Hormones were analyzed on a Q Exactive Hybrid Quadrupole-Orbitrap mass spectrometer (Thermo Scientific) using targeted selected ion monitoring in electrospray ionization mode (spray voltage, 3 kV; capillary temperature, 300°C; heater temperature, 150°C; sheath gas, 1.9 mL min⁻¹; auxiliary gas, 0.42 mL min⁻¹). Quantification was performed with embedded calibration curves using Xcalibur 2.2 SP1 and TraceFinder software. Stable isotope-labelled internal standards were included for each hormone: D6-ABA, D2-GA1, D2-GA4, D5-tZ, D3-DHZ, D6-iP, D2-IAA, D6-SA, and D2-JA (Olchemim).

### Electron and confocal microscopy

Samples were treated and observed by transmission electron microscopy (TEM) as described (Perez-Colao et al., 2025a). Yellow fluorescent protein (YFP), green fluorescent protein (GFP) and chlorophyll autofluorescence were visualized using a Zeiss 780 confocal laser scanning microscope. Excitation was performed with an argon laser at 488 nm. Emission was detected using a 450–490 nm filter for YFP or GFP and a 610–700 nm filter for chlorophyll autofluorescence. Fluorescent spots were quantified using ImageJ.

### RNA isolation and sequencing

Total RNA was extracted from approximately 4 mg of freeze-dried tissue using the PureLink™ RNA Mini Kit (Thermo Scientific), following the manufacturer’s instructions. RNA integrity was assessed using an Agilent 2100 Bioanalyzer. Samples for RNA sequencing (RNA-seq) were treated, collected, and processed as described (Perez-Colao et al., 2025a). Reads were aligned to the *A. thaliana* reference genome (GCF_000001735.4_TAIR10.1; TAIR10), with an average alignment rate of 97.93%. Gene expression quantification and identification of differentially expressed genes (DEGs) were performed using the default DrTom analysis pipeline provided by BGI. Heatmaps were created using the pheatmap package in RStudio. (Kolde, 2019). Transcripts per million (TPM) values for selected genes are listed in **Table S2**. Raw sequencing data are deposited at the Gene Expression Omnibus (GEO) under accession number GSE303753.

### Quantitative real-time PCR

First-strand cDNA synthesis was performed using either the Transcriptor First Strand cDNA Synthesis Kit (Roche) or the NZY First-Strand cDNA Synthesis Kit (NZYtech), following the respective manufacturer’s protocols. The resulting cDNA was used as a template for RT-qPCR reactions carried out in 20 μl volumes using the LightCycler 480 SYBR Green I Master Mix (Roche) on a QuantStudio 3 (Applied Biosystems) thermocycler. Gene expression was calculated using *UBC21* (At5g25760) as a reference gene. Primers used are listed in **Table S3**.

### Protein extraction and immunoblot analysis

For protein extraction, 50 mg of fresh seedlings were ground in liquid nitrogen and homogenized in 200 μl of extraction buffer (50 mM Tris-HCl pH 7.5, 150 mM NaCl, 1 mM EDTA, 5% glycerol, 0.1% SDS, 1 mM DTT, and a protease inhibitor cocktail from Roche). Homogenates were incubated on ice for 10 min and centrifuged (15,000x*g*, 10 min, 4°C). The supernatant was collected and protein concentration determined using Bradford reagent (Thermo Scientific), with BSA as standard. For Immunoblot analysis, equal amounts of protein (20 μg) were mixed with 4× Laemmli buffer (125 mM Tris-HCl pH 6.8, 4% SDS, 20% glycerol, 10% β-mercaptoethanol, 0.01% bromophenol blue), denatured at 95°C for 5 min, and resolved on 12% SDS-PAGE gels. Proteins were transferred to PVDF membranes (Immobilon-P, Millipore) at 100 V for 90 min in Towbin buffer (25 mM Tris, 192 mM glycine, 20% methanol). Membranes were blocked in 5% non-fat dried milk in TBS-T (20 mM Tris-HCl pH 7.5, 150 mM NaCl, 0.1% Tween-20) for 1 h at room temperature, and incubated overnight at 4°C with a 1:1000 dilution of anti-SAL1 (AS07256, Agrisera) in TBS-T containing 1% non-fat dried milk. After washing, membranes were incubated for 1 h at room temperature with anti-rabbit HRP-conjugated secondary antibodies (Agrisera) and visualized using an ECL detection system (GE Healthcare). In parallel, equal protein aliquots were resolved on a separate SDS-PAGE gel and stained with Coomassie Brilliant Blue.

### Cloning and plasmid construction

Coding sequences of *GLK1* (At2g20570) and *FBN7a* (At3g23400) were amplified from *A. thaliana* Col-0 cDNA using Phusion High-Fidelity DNA polymerase (Thermo Scientific) and cloned into the pDONR207 entry vector via BP recombination using Gateway™ BP Clonase™ II (Invitrogen), following the manufacturer’s instructions. Entry clones were verified by sequencing and subsequently transferred into the destination vector pGWB405 by LR recombination using Gateway™ LR Clonase™ II (Invitrogen). For recombinant protein production, the coding sequence of *SAL1* (At5g63980) lacking the chloroplast transit peptide was synthesized as a G-Block (Integrated DNA Technologies, IDT) containing the restriction sites *Nco*I and *Eco*RI, and cloned into the bacterial expression vector pETM11 by restriction–ligation cloning. All plasmids were confirmed by Sanger sequencing before transformation into expression hosts. Primer and G-Block sequences are listed in **Table S3**.

### Transient expression in *Nicotiana benthamiana*

Three-week-old *N. benthamiana* plants were agroinfiltrated with *Agrobacterium tumefaciens* strain GV3101 carrying the appropriate FBN7a-GFP or/and GLK1 constructs. For co-infiltration assays, bacterial suspensions were prepared in infiltration buffer (10 mM MES pH 5.6, 10 mM MgCl₂, 150 μM acetosyringone) and incubated at room temperature for 2–3 h before infiltration. GLK1 cultures were adjusted to OD₆₀₀ = 0.5, FBN7a-GFP cultures to OD₆₀₀ = 0.1, and an additional culture carrying the silencing suppressor HCPro (Perez-Colao et al., 2025b) was included at OD₆₀₀ = 0.1. One day after agroinfiltration, the same leaf areas were infiltrated with either 100 μM X57 or 0.1% DMSO as a mock control. Samples were analyzed by confocal microscopy 72 h post-infiltration, and harvested 4 days after infiltration (3 days after treatment) for metabolite quantification.

### Recombinant protein expression and purification

The construct pETM11-*SAL1* was transformed into *Escherichia coli* BL21 (DE3) cells. A single colony was used to inoculate LB medium containing 50 µg mL⁻¹ kanamycin and cultured overnight at 37°C. The following day, cultures were diluted 1:100 into fresh LB medium with kanamycin and grown at 37°C until OD₆₀₀ reached 0.6. Protein expression was induced with 1 mM IPTG, and cells were incubated overnight at 16°C. Cells were harvested by centrifugation (5,000 rpm, 15 min, 4°C), resuspended in lysis buffer (50 mM Tris-HCl pH 8.0, 150 mM NaCl, 20 mM KCl, 1 mM MgCl₂, 5% glycerol, 2 mM β-mercaptoethanol) and disrupted by sonication on ice. Lysates were cleared by centrifugation (20,000 rpm, 30 min, 4°C). The soluble fraction was applied to a Ni-NTA HiTrap FF column (Cytiva) equilibrated with lysis buffer. After washing with buffer containing 40 mM imidazole, the His-tagged SAL1 protein was eluted with 300 mM imidazole in the same buffer. Eluted fractions were pooled and concentrated with Amicon Ultra centrifugal filters (30 kDa cutoff, Millipore). Samples were then loaded onto a HiLoad 26/60 Superdex 200 column (Cytiva) connected to an ÄKTA Go FPLC system (Cytiva), pre-equilibrated with SEC buffer (50 mM Tris-HCl pH 8.0, 150 mM NaCl, 20 mM KCl, 1 mM MgCl₂, 2 mM β-mercaptoethanol). Protein was separated at 0.5 mL min⁻¹, and monomeric fractions were collected, concentrated by Amicon Ultra centrifugal filters, quantified by NanoDrop, and flash frozen in aliquots at −80°C. The final storage buffer consisted of 50 mM Tris-HCl pH 8.0, 150 mM NaCl, 20 mM KCl, 1 mM MgCl₂, 2 mM β-mercaptoethanol, and 15% glycerol. Protein purity was assessed by SDS-PAGE and Coomassie Brilliant Blue staining.

### Isothermal shift assay

Protein–small molecule interactions were analyzed using the isothermal shift assay (iTSA) as described (Ball et al., 2020) with minor modifications. *A. thaliana* Col-0 seedlings germinated and grown on MS plates for 10 days were homogenized in liquid nitrogen and resuspended in extraction buffer (50 mM HEPES pH 7.5, 150 mM NaCl, 1 mM EDTA, 5% glycerol, 1 mM DTT, protease inhibitor cocktail). After centrifugation (20,000 rpm, 15 min, 4°C), the soluble fraction was divided into aliquots and spiked for 30 min at 4°C with either 100 µM X57 or 100 µM X56. Samples were subjected to heat treatment at 53°C for 3 min, immediately cooled on ice, and centrifuged again (20,000 rpm, 15 min, 4°C) to remove precipitated proteins. Supernatants were collected and mixed with four volumes of cold acetone to precipitate proteins overnight at −20°C. Pellets were washed once with 80% acetone, air-dried, and resuspended in 100 mM Tris-HCl (pH 8.5) containing 4% sodium deoxycholate (SDC). Samples were reduced with 10 mM Tris (2-carboxyethyl) phosphine and alkylated with 40 mM chloroacetamide for 5 min at 45°C with shaking at 2000 rpm in an Eppendorf ThermoMixer C. Trypsin, prepared in 50 mM ammonium bicarbonate, was added at a 1:100 enzyme-to-protein ratio, and digestion was carried out overnight at 37°C with shaking (1500 rpm) in a final volume of 300 µL. After digestion, SDC was removed by phase extraction, and samples were acidified to 1% trifluoroacetic acid and desalted using C18 StageTips. Approximately 1 µL (∼200 ng) of each peptide digest was loaded onto a Thermo EASY-nLC 1200 equipped with a Thermo Acclaim PepMap RSLC 0.1 mm × 20 mm C18 trapping column, washed for ∼5 min with buffer A (0.1% formic acid in water), and subsequently separated on a Thermo Acclaim PepMap RSLC 0.075 mm × 250 mm analytical column. Chromatographic separation was performed at 300 nL min^-1^ using a 35 min gradient consisting of 5% to 19% buffer B (80% acetonitrile and 0.1% formic acid in water) from 0 to 19 min, 19% to 40% B from 19 to 24 min, and 40% to 90% B from 24 to 25 min, followed by a high-organic wash. Eluted peptides were ionized using a FlexSpray source and analyzed on a Thermo Q-Exactive HF-X mass spectrometer operating in DIA mode. Full MS scans were acquired at 45,000 resolution (m/z 200) over a 400–1000 m/z range, followed by sequential fragmentation of 30 DIA windows of 20 m/z each using HCD, with fragment spectra measured at 15,000 resolution (m/z 200). DIA data were processed using DIA-NN v1.8.1 in library-free mode with Deep Learning and the Robust LC (high-precision) quantitation strategy, employing RT-dependent cross-referencing. Spectra were searched against the *A. thaliana* proteome (UniProt, downloaded 2024-09-16) supplemented with common contaminants (cRAP, TheGPM). DIA-NN automatically optimized search parameters, and identifications were filtered at 1% precursor FDR. Differential protein enrichment between X57 and X56 conditions was assessed using Student’s t-test with a significance threshold of p < 0.05.

### Nano differential scanning fluorimetry and microscale thermophoresis

SAL1-small molecule binding was assessed using nanoDSF (Prometheus NT.48, NanoTemper) and purified recombinant His-tagged SAL1. Protein samples (2 µM SAL1 in 50 mM Tris-HCl pH 8.0, 150 mM NaCl, 20 mM KCl, 1 mM MgCl₂, 2 mM β-mercaptoethanol, 15% glycerol) were incubated with 100 µM X57, 100 µM X56, or 0.1% DMSO for 10 min at room temperature. Samples (10 µL) were loaded into standard capillaries, and intrinsic tryptophan/tyrosine fluorescence at 330 and 350 nm was recorded during a linear temperature ramp from 25°C to 90°C at 1°C min⁻¹. The SAL1 melting temperature (Tₘ) was calculated as the inflection point of the fluorescence ratio (F350/F330) using PR.ThermControl software (NanoTemper). Binding affinity between SAL1 and X57 was analyzed using microscale thermophoresis (MST) on a Monolith NT.115 (NanoTemper). Recombinant SAL1 was fluorescently labeled with the RED-tris-NTA dye (NanoTemper) following the manufacturer’s protocol. A constant concentration of labeled protein (50 nM) was mixed with a serial dilution of X57 (0.1 nM–1 mM) or X56 (negative control) in assay buffer (50 mM Tris-HCl pH 8.0, 150 mM NaCl, 20 mM KCl, 1 mM MgCl₂, 0.05% Tween-20). After 10 min incubation at room temperature, samples were loaded into premium-coated capillaries, and thermophoresis was measured at 40% MST power and 20% LED power. Data were analyzed with MO.Affinity Analysis software (NanoTemper) using the standard Kd fit model.

### *In vitro* SAL1 phosphatase activity assay

Phosphatase activity of recombinant His-tagged SAL1 was measured using *para*-methylumbelliferyl phosphate (pMUP; Sigma-Aldrich) as a fluorogenic substrate. Reactions were performed in 96-well plates in a final volume of 100 µL containing assay buffer (50 mM Tris-HCl pH 8.0, 150 mM NaCl, 20 mM KCl, 1 mM MgCl₂, 2 mM β-mercaptoethanol). Purified SAL1 protein (50 nM) was incubated with increasing concentrations of X57, X56, or 0.1% DMSO (mock) for 20 min at room temperature prior to substrate addition. Reactions were initiated by adding 100 µM pMUP. Release of fluorescent 4-methylumbelliferone (4-MU) was monitored in a microplate reader (Tecan Spark) using excitation at 360 nm and emission at 450 nm. Fluorescence was recorded every minute for up to 30 min. Enzymatic activity was expressed as Relative Fluorescence units, normalized to mock controls. Dose–response curves were fitted using nonlinear regression (four-parameter logistic model) in GraphPad Prism 9 to calculate the half-maximal inhibitory concentration (IC₅₀).

## ACKNOWLEDGEMENTS

We thank the Michigan State University Proteomics Facility and Dr. Douglas Whitten for their expert assistance in conducting the proteomic analyses, and the technical support of Hannah McKinnon-Reish, Hillary Fischer, Pallavi Agarwal, and Jieun Kang for with NanoDSF and MST. We are also grateful to the staff of the IBMCP Metabolomics Platform for assistance with HPLC and hormone analyses, Lidia Orea-Ordoñez and Rafa Ruiz-Partida for help with protein purification and enzymatic assays and José M. Pérez-Beser and Esther Pau-Ramos for general technical assistance. We acknowledge María T. Mínguez (SCSIE Microscopy Service, University of Valencia) for her expert support with electron microscopy, Felix Kessler (University of Neuchatel) for the *35S:FBN7a-YFP* line, and Dr. Jose Luis Micol (University Miguel Hernandez) for providing the seeds of the *sal1* allele *ron1-1*.

## FUNDING

This work was funded by grants from the Spanish Agencia Estatal de Investigación (AEI, MICIU/AEI/10.13039/501100011033), Generalitat Valenciana, “ERDF A way of making Europe” and “European Union NextGeneration EU/PRTR” to MR-C (grants PID2020-115810GB-I00, RED2022-134577-T, PID2023-149584NB-I00, and AGROALNEXT/2022/067), JL-L (grants RYC2020-029097-I, PID2021-128826OA-I00, CISEJI/2022/26, and CNS2023-145540) and GalChimia (grant RTC-2017-6019-2). Work was also supported by the National Institute of General Medical Sciences of the National Institutes of Health under award number R35GM153298 to AS and the German Research Foundation (DFG) under Germanýs Excellence Strategy EXC 2048/1 project 390686111 to SK and AK. PP-C received a predoctoral fellowship from the Generalitat Valenciana (CIACIF/2021/278).

## SUPPLEMENTAL FIGURES

Figure S1. X57 causes loss of photosynthetic efficiency in a reversible manner.

Figure S2. Hormone levels in seedlings treated with X57 for 5 h.

Figure S3. Phenotype of seedlings 48 h after transferring to X57-supplemented liquid medium.

Figure S4. X56 is a dead analog of X57.

Figure S5. X57 treatment does not change PAP levels in the *sal1* mutant.

Figure S6. Genes targeted by the SAL1-PAP pathway respond to X57 treatment.

Figure S7. X57 sensitivity of different retrograde signaling mutants.

## Notes

### Competing Interest Statement

The authors have declared no competing interest.

## REFERENCES

Ashykhmina, N., Lorenz, M., Frerigmann, H., Koprivova, A., Hofsetz, E., Stührwohldt, N., Flügge, U.I., Haferkamp, I., Kopriva, S., & Gigolashvili, T. (2019). PAPST2 Plays Critical Roles in Removing the Stress Signaling Molecule 3’-Phosphoadenosine 5’-Phosphate from the Cytosol and Its Subsequent Degradation in Plastids and Mitochondria. Plant Cell 31(1), 231–249. DOI: 10.1105/tpc.18.00512.

Ball, K. A., Webb, K. J., Coleman, S. J., Cozzolino, K. A., Jacobsen, J., Jones, K. R., Stowell, M. H. B., & Old, W. M. (2020). An isothermal shift assay for proteome scale drug-target identification. Communications Biology 3(1), 75. DOI: 10.1038/S42003-020-0795-6;TECHMETA

Cackett, L., Luginbuehl, L. H., Schreier, T. B., Lopez-Juez, E., & Hibberd, J. M. (2022). Chloroplast development in green plant tissues: the interplay between light, hormone, and transcriptional regulation. New Phytologist 233(5), 2000–2016. DOI: 10.1111/NPH.17839

Chan, K. X., Phua, S. Y., Crisp, P., McQuinn, R., & Pogson, B. J. (2016). Learning the Languages of the Chloroplast: Retrograde Signaling and Beyond. Annual Review of Plant Biology 67, 25–53. DOI: 10.1146/ANNUREV-ARPLANT-043015-111854

Choi, H., Yi, T., & Ha, S. H. (2021). Diversity of Plastid Types and Their Interconversions. Frontiers in Plant Science 12, 692024. DOI: 10.3389/FPLS.2021.692024/PDF

Cortleven, A., & Schmülling, T. (2015). Regulation of chloroplast development and function by cytokinin. Journal of Experimental Botany 66(16), 4999–5013. DOI: 10.1093/jxb/erv132

Estavillo, G. M., Crisp, P. A., Pornsiriwong, W., Wirtz, M., Collinge, D., Carrie, C., Giraud, E., Whelan, J., David, P., Javot, H., Brearley, C., Hell, R., Marin, E., & Pogson, B. J. (2011). Evidence for a SAL1-PAP Chloroplast Retrograde Pathway That Functions in Drought and High Light Signaling in Arabidopsis. Plant Cell 23(11), 3992–4012. DOI: 10.1105/TPC.111.091033

Gao, K., Oerlemans, R., & Groves, M. R. (2020). Theory and applications of differential scanning fluorimetry in early-stage drug discovery. Biophysical Reviews 12(1), 85–104. DOI: 10.1007/S12551-020-00619-2

García, I. B., Ledezma, A. K. D., Montaño, E. M., Leyva, J. A. S., Carrera, E., & Ruiz, I. O. (2020). Identification and Quantification of Plant Growth Regulators and Antioxidant Compounds in Aqueous Extracts of Padina durvillaei and Ulva lactuca. Agronomy 10(6), 866. DOI: 10.3390/AGRONOMY10060866

Gil, M. J., Coego, A., Mauch-Mani, B., Jordá, L., & Vera, P. (2005). The Arabidopsis csb3 mutant reveals a regulatory link between salicylic acid-mediated disease resistance and the methyl-erythritol 4-phosphate pathway. Plant Journal 44(1), 155–166. DOI: 10.1111/J.1365-313X.2005.02517.X

Hernández-Verdeja, T., & Strand, Å. (2018). Retrograde Signals Navigate the Path to Chloroplast Development. Plant Physiology, 176(2), 967–976. DOI: 10.1104/PP.17.01299

Jarvis, P., & López-Juez, E. (2013). Biogenesis and homeostasis of chloroplasts and other plastids. Nature Reviews in Molecular Cell Biology 14(12), 787–802. DOI: 10.1038/nrm3702.

Jiang, J., & Dehesh, K. (2021). Plastidial retrograde modulation of light and hormonal signaling: an odyssey. New Phytologist 230(3), 931–937. DOI: 10.1111/NPH.17192

Khin, N. C., Carmody, M., Schwartz, B. D., Dlugosch, M., Yee, S., Moore, M., Gardiner, M., Tan, L. L., Jackson, C., Corry, B., Malins, L. R., Pogson, B., & Chan, K. X. (2025). High-throughput screening and structure-guided design of small molecules enable modulation of SAL-PAP stress signaling. BioRxiv, DOI: 10.1101/2025.10.20.683270

Kolde R. (2019). Pheatmap: pretty heatmaps. R package v.1.0.12. [WWW document] URL https://CRAN.R-project.org/package=pheatmap [accessed 26 October 2025].

Larkin, R. M., Alonso, J. M., Ecker, J. R., & Chory, J. (2003). GUN4, a regulator of chlorophyll synthesis and intracellular signaling. Science 299(5608), 902–906. DOI: 10.1126/SCIENCE.1079978

Li, L., Rodríguez-Concepción, M., & Al-Babili, S. (2025). Advances in carotenoid and apocarotenoid metabolisms and functions in plants. Plant Physiology 198(4), kiaf304. DOI: 10.1093/PLPHYS/KIAF304

Li, Y., Zhou, Y., Wang, Z., Sun, X., & Tang, K. (2010). Engineering tocopherol biosynthetic pathway in Arabidopsis leaves and its effect on antioxidant metabolism. Plant Science 178(3), 312–320. DOI: 10.1016/J.PLANTSCI.2010.01.004

Llorente, B., Torres-Montilla, S., Morelli, L., Florez-Sarasa, I., Matus, J. T., Ezquerro, M., D’Andrea, L., Houhou, F., Majer, E., Picó, B., Cebolla, J., Troncoso, A., Fernie, A. R., Daròs, J. A., & Rodriguez-Concepcion, M. (2020). Synthetic conversion of leaf chloroplasts into carotenoid-rich plastids reveals mechanistic basis of natural chromoplast development. Proceedings of the National Academy of Sciences of the United States of America, 117(35), 21796–21803. DOI: 10.1073/PNAS.2004405117

Lu, Y., Rijzaani, H., Karcher, D., Ruf, S., & Bock, R. (2013). Efficient metabolic pathway engineering in transgenic tobacco and tomato plastids with synthetic multigene operons. Proceedings of the National Academy of Sciences of the United States of America 110(8), E623–32. DOI: 10.1073/PNAS.1216898110

Magnusson, A. O., Szekrenyi, A., Joosten, H. J., Finnigan, J., Charnock, S., & Fessner, W. D. (2018). nanoDSF as screening tool for enzyme libraries and biotechnology development. The Febs Journal 286(1), 184. DOI: 10.1111/FEBS.14696

Mesa, T., Mariani, C., & Munné-Bosch, S. (2025). Postharvest regreening: a species- and variety-dependent process triggered by phytohormones and light. Annals of Botany 10, mcaf094. DOI: 10.1093/aob/mcaf094

Mochizuki, N., Brusslan, J. A., Larkin, R., Nagatani, A., & Chory, J. (2001). Arabidopsis genomes uncoupled 5 (GUN5) mutant reveals the involvement of Mg-chelatase H subunit in plastid-to-nucleus signal transduction. Proceedings of the National Academy of Sciences of the United States of America 98(4), 2053. DOI: 10.1073/PNAS.98.4.2053

Myouga, F., Akiyama, K., Tomonaga, Y., Kato, A., Sato, Y., Kobayashi, M., Nagata, N., Sakurai, T., & Shinozaki, K. (2013). The Chloroplast Function Database II: a comprehensive collection of homozygous mutants and their phenotypic/genotypic traits for nuclear-encoded chloroplast proteins. Plant Cell and Physiology 54(2), e2. DOI: 10.1093/pcp/pcs171

Ortiz-Alcaide, M., Llamas, E., Gomez-Cadenas, A., Nagatani, A., Martínez-García, J. F., & Rodríguez-Concepción, M. (2019). Chloroplasts modulate elongation responses to canopy shade by retrograde pathways involving hy5 and abscisic acid. Plant Cell 31(2), 384–398. DOI: 10.1105/TPC.18.00617

Perez-Colao, P., Glauser, G., Cruces, J., Lozano-Juste, J., & Rodriguez-Concepción, M. (2025a). A chemical probe for increasing leaf tocopherol levels by coordinated modulation of biosynthesis, competition, and storage. Plant Biotechnology Journal, DOI:10.1111/pbi.70459.

Perez-Colao, P., Morelli, L., & Rodriguez-Concepcion, M. (2025b). Using *Agrobacterium tumefaciens* to Assemble Multi-step Metabolic Pathways in *Nicotiana benthamiana*. Methods in Molecular Biology 2911, 11–20. DOI: 10.1007/978-1-0716-4450-8_3.

Phua, S. Y., Yan, D., Chan, K. X., Estavillo, G. M., Nambara, E., & Pogson, B. J. (2018). The arabidopsis SAL1-PAP pathway: A case study for integrating chloroplast retrograde, light and hormonal signaling in modulating plant growth and development? Frontiers in Plant Science 9, 1171. DOI: 10.3389/FPLS.2018.01171/FULL

Ramel, F., Birtic, S., Ginies, C., Soubigou-Taconnat, L., Triantaphylidès, C., & Havaux, M. (2012). Carotenoid oxidation products are stress signals that mediate gene responses to singlet oxygen in plants. Proceedings of the National Academy of Sciences of the United States of America 109(14), 5535–5540. DOI: 10.1073/PNAS.1115982109

Robles, P., Fleury, D., Candela, H., Cnops, G., Alonso-Peral, M. M., Anami, S., Falcone, A., Caldana, C., Willmitzer, L., Ponce, M. R., van Lijsebettens, M., & Micol, J. L. (2009). The RON1/FRY1/SAL1 Gene Is Required for Leaf Morphogenesis and Venation Patterning in Arabidopsis. Plant Physiology 152(3), 1357. DOI: 10.1104/PP.109.149369

Rossini, L., Cribb, L., Martin, D. J., & Langdale, J. A. (2001). The maize golden2 gene defines a novel class of transcriptional regulators in plants. Plant Cell 13(5), 1231–1244. DOI: 10.1105/TPC.13.5.1231

Sauret-Güeto, S., Botella-Pavía, P., Flores-Pérez, U., Martínez-García, J. F., San Román, C., León, P., Boronat, A., & Rodríguez-Concepción, M. (2006). Plastid cues posttranscriptionally regulate the accumulation of key enzymes of the methylerythritol phosphate pathway in Arabidopsis. Plant Physiology 141(1), 75–84. DOI: 10.1104/PP.106.079855

Schlossarek, D., Luzarowski, M., Sokołowska, E. M., Thirumalaikumar, V. P., Dengler, L., Willmitzer, L., Ewald, J. C., & Skirycz, A. (2022). Rewiring of the protein-protein-metabolite interactome during the diauxic shift in yeast. Cellular and Molecular Life Sciences 79(11). DOI: 10.1007/S00018-022-04569-8

Seidel, S. A. I., Dijkman, P. M., Lea, W. A., van den Bogaart, G., Jerabek-Willemsen, M., Lazic, A., Joseph, J. S., Srinivasan, P., Baaske, P., Simeonov, A., Katritch, I., Melo, F. A., Ladbury, J. E., Schreiber, G., Watts, A., Braun, D., & Duhr, S. (2013). Microscale thermophoresis quantifies biomolecular interactions under previously challenging conditions. Methods 59(3), 301–315. DOI: 10.1016/j.ymeth.2012.12.005

Sierra, J., Escobar-Tovar, L., & Leon, P. (2023). Plastids: diving into their diversity, their functions, and their role in plant development. Journal of Experimental Botany 74(8), 2508–2526. DOI: 10.1093/jxb/erad044

Susek, R. E., Ausubel, F. M., & Chory, J. (1993). Signal transduction mutants of Arabidopsis uncouple nuclear CAB and RBCS gene expression from chloroplast development. Cell 74(5), 787–799. DOI: 10.1016/0092-8674(93)90459-4

Vidi, P. A., Kessler, F., & Bréhélin, C. (2007). Plastoglobules: A new address for targeting recombinant proteins in the chloroplast. BMC Biotechnology 7, 4. DOI: 10.1186/1472-6750-7-4,

Waters, M. T., Wang, P., Korkaric, M., Capper, R. G., Saunders, N. J., & Langdale, J. A. (2009). GLK Transcription Factors Coordinate Expression of the Photosynthetic Apparatus in Arabidopsis. Plant Cell 21(4), 1109–1128. DOI: 10.1105/TPC.108.065250

Xiao, Y., Savchenko, T., Baidoo, E. E. K., Chehab, W. E., Hayden, D. M., Tolstikov, V., Corwin, J. A., Kliebenstein, D. J., Keasling, J. D., & Dehesh, K. (2012). Retrograde signaling by the plastidial metabolite MEcPP regulates expression of nuclear stress-response genes. Cell 149(7), 1525–1535. DOI: 10.1016/j.cell.2012.04.038

Zhao C, Wang Y, Chan KX, Marchant DB, Franks PJ, Randall D, Tee EE, Chen G, Ramesh S, Phua SY, Zhang B, Hills A, Dai F, Xue D, Gilliham M, Tyerman S, Nevo E, Wu F, Zhang G, Wong GK, Leebens-Mack JH, Melkonian M, Blatt MR, Soltis PS, Soltis DE, Pogson BJ, & Chen ZH (2019). Evolution of chloroplast retrograde signaling facilitates green plant adaptation to land. Proceedings of the National Academy of Sciences of the United States of America 116(11), 5015–5020. DOI: 10.1073/pnas.1812092116.

