## Supplemental figures for "Chemical modulation of chloroplast de- and re-differentiation reveals a role for the SAL1-PAP retrograde pathway in facilitating plastid transitions"

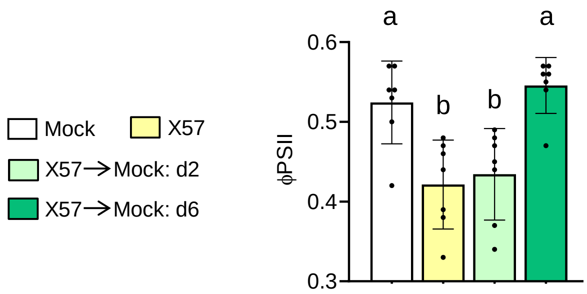


**Figure S1. X57 causes loss of photosynthetic efficiency in a reversible manner.** Wild-type *Arabidopsis thaliana* seedlings were germinated and grown for 5 days on plates containing 0.05% DMSO and then transferred to new plates supplemented with either 0.05% DMSO (mock) or 40 µM X57 for 5 more days. Some X57-treated seedlings were subsequently transferred to mock medium and grown for two (d2) or six (d6) additional days. Effective quantum yield of photosystem II (ΦPSII) was measured on true leaves of several seedlings grown on different plates for each treatment. Mean and SD of seven biological replicates (n=7) are shown. Statistically significant differences are represented with letters (one-way ANOVA followed by Tukey's post hoc test, $p$<0.05).

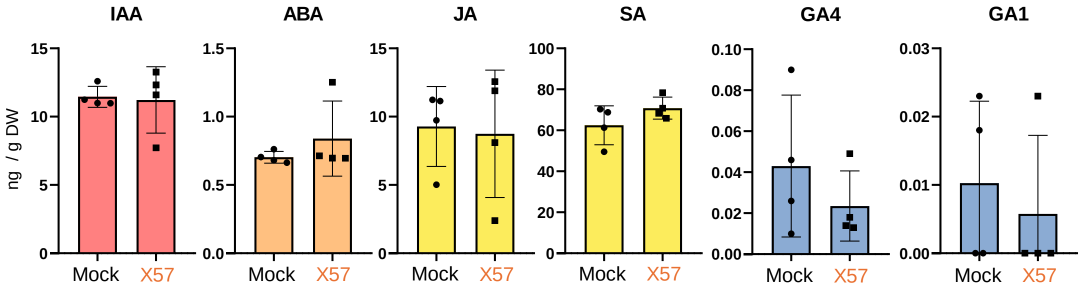


**Figure S2. Hormone levels in seedlings treated with X57 for 5 h.** Levels of the indicated hormones in *A. thaliana* WT seedlings germinated and grown for 5 days on plates containing 0.05% DMSO and then transferred to liquid medium supplemented with either 0.05% DMSO (mock) or 100 µM X57 for 5 h. Mean and SD of four independent samples (n=4) are represented. No statistically significant differences relative to mock samples (t-test, $p$<0.05) were found for any of the hormones. DW, dry weight. IAA, indole-3-acetic acid; ABA, abscisic acid; JA, jasmonic acid; SA, salicylic acid; GA4, gibberellin $A_4$; GA1, gibberellin $A_1$.

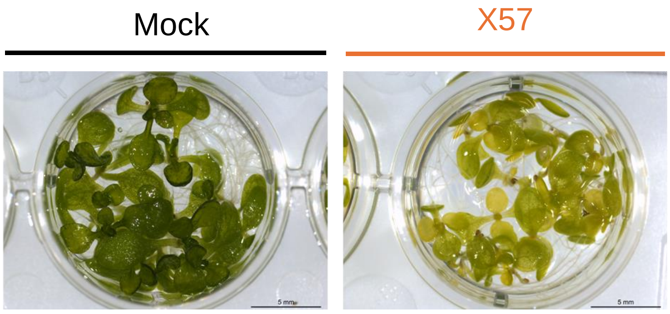


**Figure S3. Phenotype of seedlings 48 h after transferring to X57-supplemented liquid medium.** *A. thaliana* WT seedlings germinated and grown for 5 days in mock plates were transferred to fresh liquid medium supplemented with 100 µM X57 or 0.05% DMSO (mock). Images were taken 48 h after transfer.

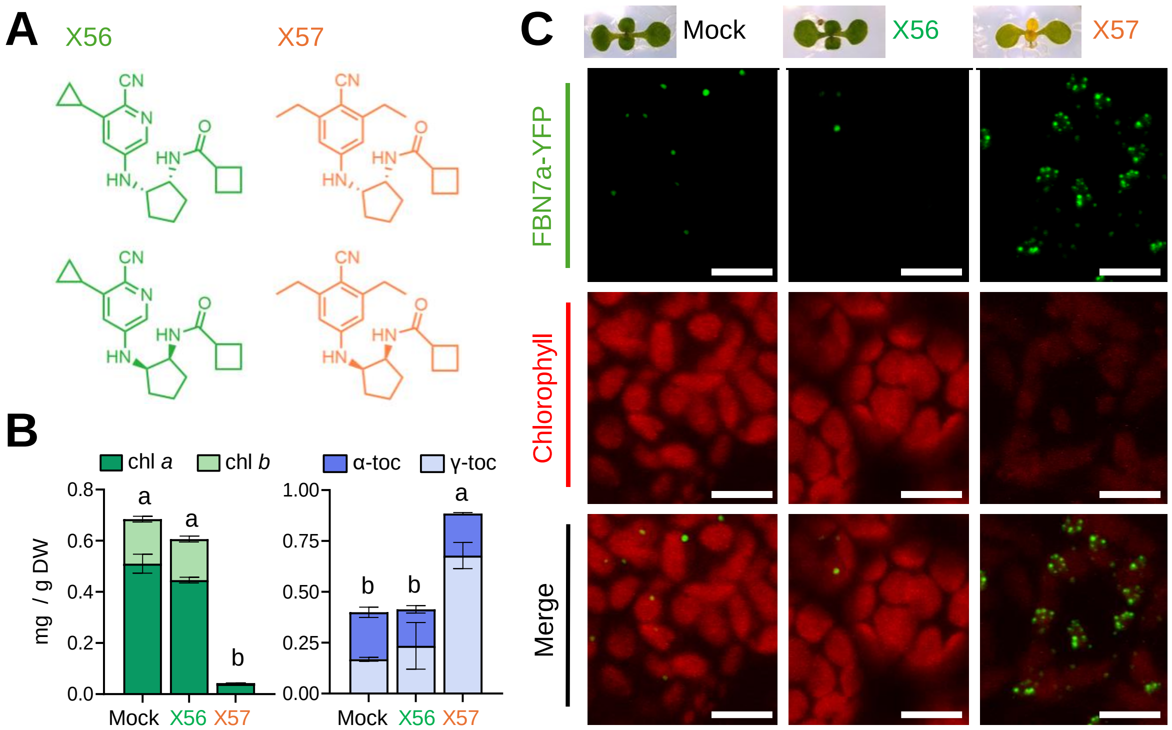


**Figure S4. X56 is a dead analog of X57.** (A) Chemical structures of X56 and X57. The two diastereomers of each compound represented in the picture were present in the mix used for experiments. (B) Levels of chlorophylls (*a* and *b*) and tocopherols (α and γ) in WT seedlings germinated and grown for 5 days in mock medium and then transferred for 3 days to fresh liquid medium supplemented with 0.05% DMSO (mock), 100 µM X57, or 100 µM X56. Mean and SD of three independent samples (n=3) are represented. Statistically significant differences are represented with letters (one-way ANOVA followed by Tukey's post hoc test, $p$<0.05) of total chlorophyll or tocopherol contents. DW, dry weight. (C) Confocal microscopy analysis of PG distribution in the leaves of transgenic *35S:FBN7a-YFP* seedlings grown as described in (B). Panels show fluorescence from the PG marker protein FBN7a-YFP (in green), chlorophyll autofluorescence (in red), or both, in the same microscopy field. Bars, 5 µm.

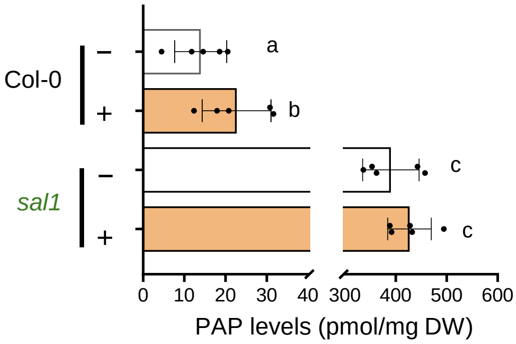


**Figure S5. X57 treatment does not change PAP levels in the *sal1* mutant.** PAP contents in *A. thaliana* WT (Col-0) and mutant *sal1* seedlings germinated and grown for 5 days in medium supplemented with 0.05% DMSO (-) or 25 µM X57 (+). Mean and SD of n=5 independent biological replicates are shown. Letters indicate statistically significant differences ($p$<0.05) according to one-way ANOVA followed by Tukey's post hoc test. DW, dry weight.

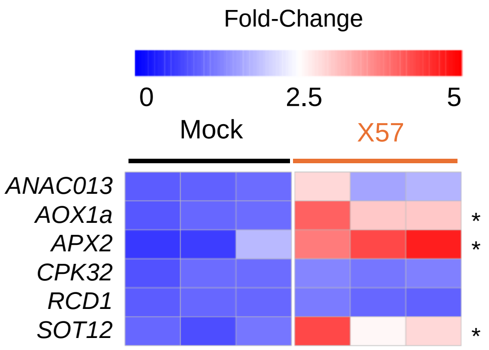


**Figure S6. Genes targeted by the SAL1-PAP pathway respond to X57 treatment.** *A. thaliana* WT seedlings were germinated and grown for 5 days on plates containing 0.05% DMSO and then transferred to liquid medium supplemented with either 0.05% DMSO (mock) or 100 µM X57. Samples for RNA-seq were collected 5 h after transfer. Heatmaps shows transcript abundance as Fold-Change relative to mock samples. Columns correspond to biological replicates. Asterisks indicate significant changes in X57 compared to mock samples (DESeq2; FDR ≤ 0.05). Gene accessions and TPM values are listed in Table S2.

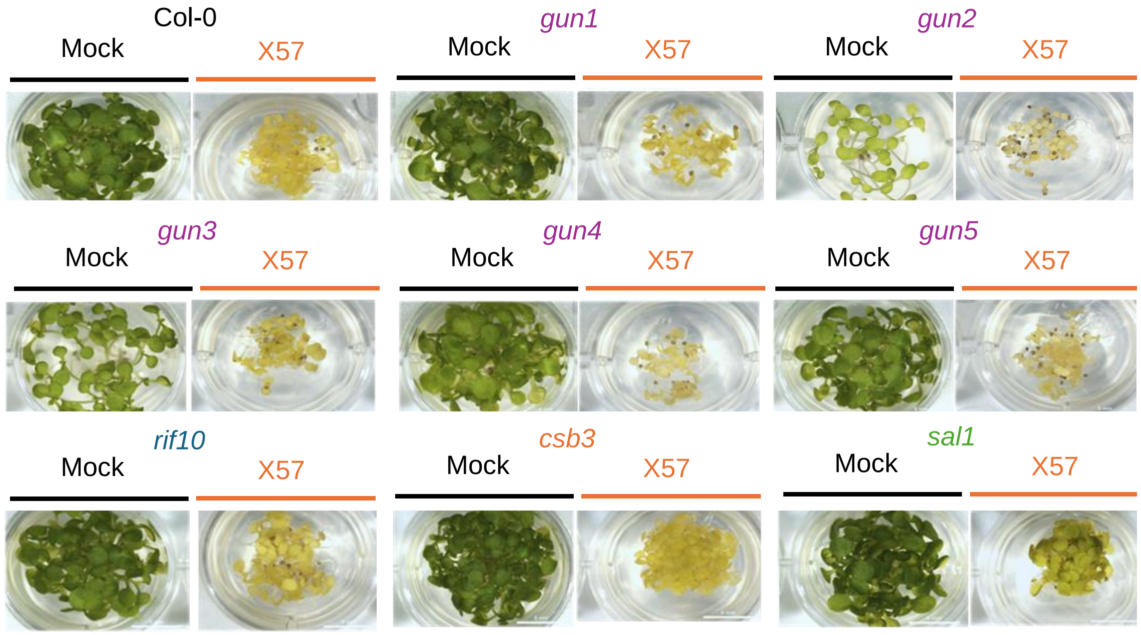


**Figure S7. X57 sensitivity of different retrograde signaling mutants.** Representative images *A. thaliana* WT (Col-0) and retrograde signaling mutant seedlings germinated and grown for 7 days in solid medium supplemented with 0.05% DMSO (mock) or 25 µM X57.

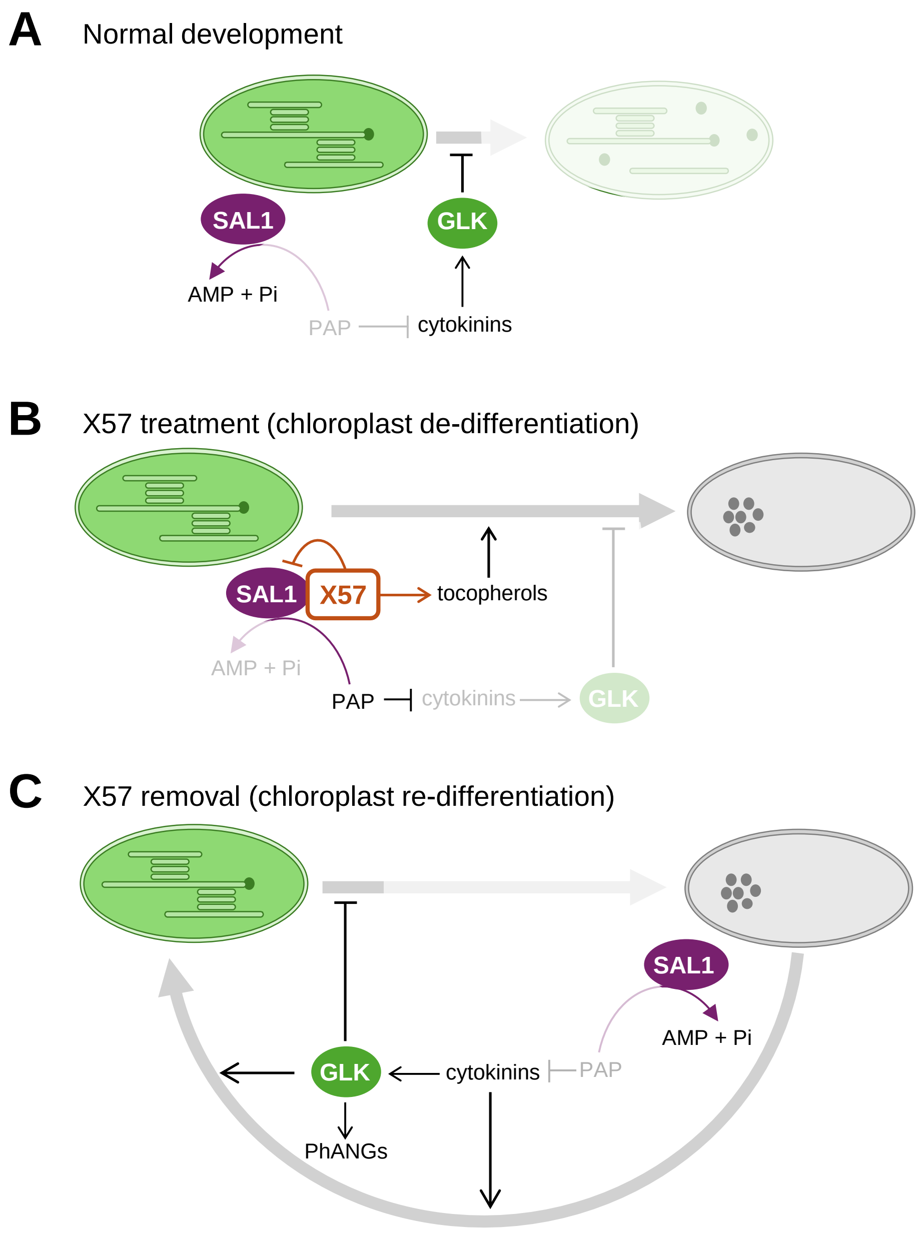


**Figure S8. Model of X57-dependent chloroplast de- and re-differentiation.** (A) Under normal growth conditions, SAL1 degrades PAP into AMP and Pi, thereby maintaining proper cytokinin levels and *GLK1* expression to prevent loss of chloroplast identity (e.g., photosynthetic function). (B) In the presence of X57, SAL1 activity is directly inhibited, leading to the accumulation of PAP. As a consequence, cytokinin levels and *GLK1* expression are reduced and chloroplast identity declines, allowing conversion to PG-enriched plastids thanks to the SAL1-independent X57-driven induction of tocopherol accumulation. (C) Upon X57 removal, SAL1 activity is restored, allowing PAP levels and cytokinin signaling to return to normal. Reactivation of *GLK1* and downstream PhANGs expression promotes the re-differentiation of chloroplasts.
