## Supplemental table S1 for "Chemical modulation of chloroplast de- and re-differentiation reveals a role for the SAL1-PAP retrograde pathway in facilitating plastid transitions"

**Table S1.** List of proteins showing differential abundance between X57 and X56 treatments.

| | | | X57 vs X56 | | | | | | | | | | | | |
|---|---|---|---|---|---|---|---|---|---|---|---|---|---|---|---|
| **Protein** | **Locus** | **Gene Ontology (biological process)** | pvalue | FC | **LOG10Pv** | **LOG2FC** | **Mock_1** | **Mock_2** | **Mock_3** | **X57_1** | **X57_2** | **X57_3** | **X56_1** | **X56_2** | **X56_3** |
| **AT9943_LOCUS5762;AXX17_At1g72230;At1g77670** | | | 1.2595E-05 | 1.1233004 | **4.8998062** | **0.1677438** | 1.68E+06 | 1.56E+06 | 2.05E+06 | 1.73E+06 | 1.89E+06 | 1.76E+06 | 1.53E+06 | 1.69E+06 | 1.56E+06 |
| **MAP2B** | At3g59990 | protein processing [GO:0016485] | 7.9718E-05 | 1.2784606 | **4.09844388** | **0.3544077** | 960714 | 685516 | 1.36E+06 | 1.25E+06 | 1.30E+06 | 1.31E+06 | 974426 | 1.02E+06 | 1.03E+06 |
| **RPS13A** | At3g60770 | translation [GO:0006412] | 0.00017848 | 0.94456437 | **3.74842193** | **-0.08227898** | 4.13E+07 | 3.84E+07 | 4.61E+07 | 3.94E+07 | 3.68E+07 | 5.38E+07 | 4.19E+07 | 3.93E+07 | 5.64E+07 |
| **At3g53970** | At3g53970 | cellular response to auxin stimulus | 0.00023045 | 0.93410573 | **3.63742446** | **-0.09834224** | 5.12E+06 | 5.59E+06 | 3.05E+06 | 5.97E+06 | 6.25E+06 | 4.70E+06 | 6.38E+06 | 6.64E+06 | 5.11E+06 |
| **RPS3B** | At3g53870 | translation [GO:0006412] | 0.00093881 | 0.62446358 | **3.02742164** | **-0.67931066** | 3.69E+06 | 2.93E+06 | 3.32E+06 | 3.42E+06 | 2.29E+06 | 2.95E+06 | 5.05E+06 | 4.11E+06 | 4.69E+06 |
| **NIT2** | At3g44300 | indoleacetic acid biosynthetic proc | 0.00111823 | 0.99061895 | **2.95147021** | **-0.01359788** | 9.83E+06 | 8.08E+06 | 3.77E+06 | 8.45E+06 | 6.83E+06 | 4.92E+06 | 8.51E+06 | 6.90E+06 | 4.98E+06 |
| **NTF2** | At5g60980 | regulatory ncRNA-mediated gene s | 0.0013391 | 0.92712177 | **2.8731862** | **-0.10916926** | 4.70E+06 | 4.83E+06 | 2.58E+06 | 3.97E+06 | 4.52E+06 | 4.48E+06 | 4.33E+06 | 4.87E+06 | 4.79E+06 |
| **BOB1** | At5g53400 | developmental process [GO:00325( | 0.00151144 | 0.95455076 | **2.82060791** | **-0.06710618** | 2.89E+06 | 2.74E+06 | 827101 | 2.85E+06 | 2.88E+06 | 2.38E+06 | 2.97E+06 | 3.01E+06 | 2.50E+06 |
| **HSP90-7** | At4g24190 | protein secretion [GO:0009306]; re | 0.00211393 | 0.91731305 | **2.67490876** | **-0.12451393** | 1.04E+07 | 1.08E+07 | 9.00E+06 | 9.69E+06 | 1.10E+07 | 9.94E+06 | 1.06E+07 | 1.20E+07 | 1.08E+07 |
| **AXX17_At3g49250;At3g54760;T5N23_120** | | | 0.0022498 | 0.83604596 | **2.64785632** | **-0.25834584** | 470234 | 444568 | 696802 | 380043 | 340701 | 521734 | 453829 | 427435 | 604872 |
| **FACT** | At5g63560 | alkyl caffeate ester biosynthetic pro | 0.00274805 | 2.08957155 | **2.56097607** | **1.06320716** | 1.59E+06 | 1.18E+06 | 948144 | 2.75E+06 | 2.49E+06 | 2.56E+06 | 1.26E+06 | 1.25E+06 | 1.22E+06 |
| **PUX11** | At2g43210 | | 0.00288008 | 0.32198346 | **2.54059488** | **-1.63494152** | 28446.1 | 217950 | 0 | 0 | 92776.7 | 0 | 69245.3 | 160740 | 58155.9 |
| **WPP1** | At5g43070 | lateral root development [GO:0048 | 0.00372488 | 1.18043212 | **2.42888821** | **0.23931508** | 341029 | 337208 | 0 | 350204 | 384397 | 340173 | 302004 | 327546 | 280942 |
| **At2g05830** | At2g05830 | L-methionine salvage from methylt | 0.00386693 | 3.98288643 | **2.41263406** | **1.99381434** | 0 | 59565.9 | 1.16E+06 | 154348 | 292686 | 170871 | 0 | 155140 | 0 |
| **NIT4** | At5g22300 | cyanide metabolic process [GO:001 | 0.00401801 | 0.85460336 | **2.39598906** | **-0.22667311** | 1.98E+06 | 2.21E+06 | 0 | 2.04E+06 | 1.81E+06 | 838284 | 2.27E+06 | 2.10E+06 | 1.11E+06 |
| **CCT8** | At3g03960 | positive regulation of transport [GC | 0.00411072 | 0.88499688 | **2.38608188** | **-0.17625573** | 4.96E+06 | 4.35E+06 | 3.73E+06 | 4.23E+06 | 4.16E+06 | 4.32E+06 | 4.77E+06 | 4.66E+06 | 4.94E+06 |
| **P5CSB** | At3g55610 | embryo development ending in see | 0.00467207 | 0.89007889 | **2.33049112** | **-0.16799489** | 1.09E+06 | 892173 | 577748 | 872656 | 866748 | 893355 | 977859 | 964135 | 1.02E+06 |
| **TUBB6** | At5g12250 | microtubule-based process [GO:00( | 0.00478173 | 0.7580998 | **2.32041486** | **-0.39954031** | 1.61E+06 | 1.31E+06 | 1.57E+06 | 1.12E+06 | 1.19E+06 | 1.52E+06 | 1.47E+06 | 1.59E+06 | 1.97E+06 |
| **RPL32B** | At5g46430 | translation [GO:0006412] | 0.00539213 | 1.17030168 | **2.26823975** | **0.22688047** | 2.49E+06 | 2.19E+06 | 3.47E+06 | 2.96E+06 | 2.56E+06 | 3.63E+06 | 2.45E+06 | 2.16E+06 | 3.21E+06 |
| **GH3.15** | At5g13370 | auxin conjugate metabolic process | 0.00550024 | 1.41240582 | **2.25961833** | **0.49815467** | 399170 | 503528 | 271840 | 747071 | 700770 | 715291 | 566332 | 466875 | 498316 |
| **ACLB-1** | At3g06650 | acetyl-CoA biosynthetic process [G( | 0.00616942 | 0.93872543 | **2.20975536** | **-0.09122485** | 5.92E+06 | 4.41E+06 | 3.81E+06 | 4.49E+06 | 4.26E+06 | 4.85E+06 | 4.82E+06 | 4.51E+06 | 5.16E+06 |
| **PUX3** | At4g22150 | | 0.00639245 | 1.19952676 | **2.19433234** | **0.26246534** | 1.26E+06 | 1.35E+06 | 0 | 1.25E+06 | 1.46E+06 | 1.16E+06 | 1.02E+06 | 1.22E+06 | 980597 |
| **AT9943_LOCUS15222;At4g10750;T12H20.10** | | | 0.00650104 | 0 | **2.18701738** | **#¡NUM!** | 276436 | 416501 | 601281 | 0 | 0 | 0 | 311316 | 358300 | 412470 |
| **VHA-B2** | At4g38510 | ATP metabolic process [GO:004603 | 0.00714247 | 0.93394207 | **2.1461515** | **-0.09859502** | 1.75E+07 | 1.73E+07 | 1.81E+07 | 1.55E+07 | 1.51E+07 | 1.96E+07 | 1.66E+07 | 1.64E+07 | 2.06E+07 |
| **AT9943_LOCUS5170;AXX17_At1g64950;F15H11.2** | | | 0.0072175 | 1.08168172 | **2.14161326** | **0.11327606** | 692815 | 666961 | 523219 | 649568 | 681225 | 858295 | 585701 | 627532 | 810549 |
| **AXX17_At4g20330;At4g17260;LDH1;dl4665w** | | | 0.00726028 | 1.24222099 | **2.13904634** | **0.31292185** | 1.14E+06 | 710132 | 927261 | 1.04E+06 | 1.04E+06 | 1.16E+06 | 835506 | 799600 | 979300 |
| **AT9943_LOCUS21925;At5g51840;MIO24.2** | | | 0.00731577 | 1.16850949 | **2.13573985** | **0.22466946** | 448841 | 468634 | 0 | 651869 | 566932 | 532013 | 569435 | 469460 | 459436 |
| **PAP12** | At2g27190 | cellular response to phosphate star | 0.00745365 | 0.81318785 | **2.1276312** | **-0.29833944** | 1.24E+06 | 1.22E+06 | 1.16E+06 | 1.13E+06 | 1.14E+06 | 1.02E+06 | 1.36E+06 | 1.37E+06 | 1.31E+06 |
| **UGD2** | At3g29360 | carbohydrate metabolic process [G | 0.00761374 | 0.85252072 | **2.11840171** | **-0.2301932** | 5.55E+06 | 4.94E+06 | 4.52E+06 | 4.04E+06 | 4.85E+06 | 4.67E+06 | 4.92E+06 | 5.66E+06 | 5.32E+06 |
| **GAPCP1** | At1g79530 | anther wall tapetum development | 0.007983 | 0.86982692 | **2.09783383** | **-0.20119974** | 6.71E+06 | 4.81E+06 | 7.11E+06 | 5.22E+06 | 5.25E+06 | 4.92E+06 | 5.90E+06 | 5.97E+06 | 5.82E+06 |
| **AGT2** | At4g39660 | photorespiration [GO:0009853] | 0.00837283 | 1.12916996 | **2.07712766** | **0.17526265** | 8.03E+06 | 9.14E+06 | 6.38E+06 | 1.02E+07 | 9.39E+06 | 9.26E+06 | 9.18E+06 | 8.40E+06 | 7.96E+06 |
| **RGGC** | At5g47210 | regulation of translation [GO:00064 | 0.00839005 | 1.08375805 | **2.07623566** | **0.11604271** | 1.06E+07 | 1.34E+07 | 1.09E+07 | 1.17E+07 | 1.51E+07 | 1.42E+07 | 1.08E+07 | 1.41E+07 | 1.29E+07 |
| **GGP3** | At4g30550 | glucosinolate metabolic process [G | 0.008707 | 1.50519999 | **2.0601316** | **0.58995518** | 748003 | 732625 | 320457 | 941191 | 888745 | 894628 | 618504 | 639971 | 551626 |
| **AN1_LOCUS12168;AT9943_LOCUS10458;AXX17_At3g08310;At3g0** | | | 0.00879046 | 0.91802083 | **2.05598837** | **-0.12340121** | 3.11E+06 | 2.71E+06 | 2.72E+06 | 2.72E+06 | 2.39E+06 | 2.84E+06 | 2.99E+06 | 2.58E+06 | 3.09E+06 |
| **At4g10320** | At4g10320 | isoleucyl-tRNA aminoacylation [GO | 0.00880401 | 0.86491837 | **2.05531938** | **-0.20936412** | 3.18E+06 | 2.73E+06 | 3.68E+06 | 2.77E+06 | 2.49E+06 | 2.88E+06 | 3.16E+06 | 2.99E+06 | 3.26E+06 |
| **CLMP1** | At1g62390 | | 0.00907709 | 0.83935043 | **2.04205349** | **-0.25265482** | 1.65E+06 | 1.27E+06 | 1.30E+06 | 1.11E+06 | 1.27E+06 | 1.26E+06 | 1.37E+06 | 1.46E+06 | 1.49E+06 |
| **PER32** | At3g32980 | hydrogen peroxide catabolic proces | 0.00937724 | 0.8609883 | **2.0279249** | **-0.21593447** | 1.41E+07 | 1.78E+07 | 1.51E+07 | 1.63E+07 | 1.60E+07 | 1.46E+07 | 1.88E+07 | 1.89E+07 | 1.66E+07 |
| **AT9943_LOCUS11328;AXX17_At3g18820;At3g17770** | | | 0.00945451 | 0.93111865 | **2.02436108** | **-0.10296308** | 518027 | 439503 | 387203 | 573544 | 604961 | 459207 | 611617 | 639950 | 507298 |
| **PIP2-2** | At2g37170 | response to water deprivation [GO: | 0.00951244 | 0.40745653 | **2.02170819** | **-1.29528195** | 435140 | 257100 | 0 | 0 | 247742 | 365487 | 348209 | 544260 | 612548 |
| **HIPP06** | At5g03380 | cellular response to hypoxia [GO:0( | 0.0099814 | 2.67320102 | **2.00080832** | **1.41856833** | 1.74E+06 | 2.05E+06 | 296345 | 1.76E+06 | 2.03E+06 | 1.26E+06 | 848938 | 1.04E+06 | 0 |
| **PATL2** | At1g22530 | cell division [GO:0051301]; cellular | 0.01004578 | 0.85278334 | **1.99801649** | **-0.22974884** | 2.01E+06 | 1.86E+06 | 1.34E+06 | 1.76E+06 | 1.54E+06 | 1.47E+06 | 2.08E+06 | 1.76E+06 | 1.75E+06 |
| **FMT** | At3g52140 | chloroplast localization [GO:001975 | 0.01052755 | 0.84081638 | **1.97767279** | **-0.25013732** | 1.20E+06 | 1.09E+06 | 881591 | 1.09E+06 | 981066 | 738104 | 1.24E+06 | 1.14E+06 | 951693 |
| **VLN3** | At3g57410 | actin filament bundle assembly [GC | 0.0107152 | 0.96578368 | **1.96999967** | **-0.05022801** | 5.32E+06 | 5.18E+06 | 4.50E+06 | 5.25E+06 | 5.26E+06 | 4.87E+06 | 5.40E+06 | 5.46E+06 | 5.07E+06 |
| **GRIP** | At5g66030 | Golgi vesicle transport [GO:004819 | 0.01096393 | 0.73282807 | **1.96003374** | **-0.44845334** | 305220 | 355924 | 0 | 273805 | 275138 | 225753 | 349540 | 371993 | 335599 |
| **TIF3H1** | At1g10840 | abscisic acid-activated signaling pat | 0.01136914 | 0.93004105 | **1.94427257** | **-0.1046337** | 3.43E+06 | 3.74E+06 | 3.13E+06 | 3.41E+06 | 3.38E+06 | 3.32E+06 | 3.68E+06 | 3.67E+06 | 3.52E+06 |
| **At1g04170;EIF2** | | | 0.012019 | 0.91378813 | **1.9201318** | **-0.1300684** | 3.57E+06 | 3.90E+06 | 4.11E+06 | 4.01E+06 | 4.09E+06 | 4.15E+06 | 4.48E+06 | 4.43E+06 | 4.49E+06 |
| **At1g08110** | At1g08110 | | 0.01236802 | 0.8037966 | **1.90769969** | **-0.31509763** | 3.69E+06 | 3.94E+06 | 2.36E+06 | 2.94E+06 | 3.61E+06 | 3.07E+06 | 3.58E+06 | 4.38E+06 | 4.01E+06 |
| **MOS4** | At3g18165 | defense response to bacterium [GC | 0.01245558 | 0 | **1.90463593** | **#¡NUM!** | 1.15E+06 | 2.35E+06 | 264433 | 0 | 0 | 0 | 1.32E+06 | 1.84E+06 | 1.95E+06 |
| **AT9943_LOCUS18902;AXX17_At5g11310;At5g11580;C24_LOCUS21** | | | 0.01272436 | 0 | **1.89536397** | **#¡NUM!** | 428881 | 390641 | 645165 | 0 | 0 | 0 | 571417 | 515286 | 746388 |
| **GLDP1** | At4g33010 | glycine catabolic process [GO:0006 | 0.0127616 | 0.82154901 | **1.894095** | **-0.28358146** | 2.03E+07 | 2.00E+07 | 1.92E+07 | 1.85E+07 | 1.95E+07 | 1.93E+07 | 2.33E+07 | 2.39E+07 | 2.25E+07 |
| **At3g06310;At5g18800** | | | 0.0130788 | 1.22725726 | **1.88343201** | **0.2954377** | 248631 | 400231 | 437136 | 335311 | 392873 | 314063 | 261273 | 323647 | 264329 |
| **RPS20** | At3g15190 | translation [GO:0006412] | 0.01323852 | 1.1045145 | **1.87816055** | **0.14341235** | 1.03E+07 | 1.18E+07 | 4.62E+06 | 1.02E+07 | 1.20E+07 | 1.09E+07 | 9.33E+06 | 1.10E+07 | 9.66E+06 |
| **SRP19** | At1g48160 | SRP-dependent cotranslational pro | 0.01334554 | 0.52596903 | **1.87466379** | **-0.92695025** | 195214 | 184982 | 0 | 130682 | 128053 | 0 | 201989 | 194299 | 95632.6 |
| **AOC1** | At3g25760 | jasmonic acid biosynthetic process | 0.01349856 | 1.19334016 | **1.86971253** | **0.25500534** | 2.51E+06 | 2.94E+06 | 3.59E+06 | 2.80E+06 | 3.61E+06 | 2.98E+06 | 2.41E+06 | 3.05E+06 | 2.41E+06 |
| **CCH** | At3g56240 | copper ion transport [GO:0006825] | 0.01367273 | 0.72192595 | **1.86414468** | **-0.47007722** | 3.78E+07 | 3.79E+07 | 1.09E+07 | 4.24E+07 | 3.57E+07 | 2.82E+07 | 5.29E+07 | 5.16E+07 | 4.27E+07 |
| **GSTU5** | At2g29450 | glutathione metabolic process [GO: | 0.01384753 | 0.85673228 | **1.85862773** | **-0.22308364** | 6.37E+06 | 5.82E+06 | 5.52E+06 | 5.54E+06 | 5.37E+06 | 6.39E+06 | 6.69E+06 | 6.12E+06 | 7.39E+06 |
| **RPL14B** | At4g27090 | translation [GO:0006412] | 0.01415186 | 0.93030994 | **1.84918663** | **-0.10421666** | 1.29E+07 | 1.05E+07 | 1.04E+07 | 1.04E+07 | 1.09E+07 | 1.45E+07 | 1.15E+07 | 1.17E+07 | 1.53E+07 |
| **ASP2** | At5g19550 | 2-oxoglutarate metabolic process [( | 0.01428462 | 0.87793605 | **1.84513134** | **-0.18781224** | 2.29E+07 | 2.52E+07 | 1.59E+07 | 2.24E+07 | 2.35E+07 | 1.78E+07 | 2.52E+07 | 2.71E+07 | 2.03E+07 |
| **BIP3** | At1g09080 | pollen tube growth [GO:0009860]; | 0.01563931 | 0.93556053 | **1.80578239** | **-0.0960971** | 2.58E+06 | 3.63E+06 | 1.86E+06 | 2.14E+06 | 3.33E+06 | 1.96E+06 | 2.35E+06 | 3.49E+06 | 2.10E+06 |
| **MLP423** | At1g24020 | abscisic acid-activated signaling pat | 0.01622097 | 1.12817443 | **1.78992319** | **0.17399014** | 1.16E+07 | 1.13E+07 | 1.16E+07 | 1.28E+07 | 1.26E+07 | 1.38E+07 | 1.16E+07 | 1.12E+07 | 1.20E+07 |
| **LFNR2** | At1g20020 | defense response to fungus [GO:0( | 0.01671209 | 0.77143531 | **1.77696925** | **-0.37438291** | 5.77E+06 | 6.11E+06 | 3.92E+06 | 4.06E+06 | 5.71E+06 | 5.34E+06 | 5.69E+06 | 6.81E+06 | 7.08E+06 |
| **At5g64130;MHJ24.11** | | | 0.01713317 | 0.72687118 | **1.76616232** | **-0.46022839** | 2.44E+06 | 2.52E+06 | 2.20E+06 | 3.16E+06 | 3.06E+06 | 1.62E+06 | 4.08E+06 | 4.29E+06 | 2.41E+06 |
| **GSTU13** | At1g27130 | defense response by callose deposi | 0.01721904 | 0.85259354 | **1.76399116** | **-0.23006996** | 6.25E+06 | 5.96E+06 | 6.00E+06 | 6.31E+06 | 6.01E+06 | 4.55E+06 | 7.25E+06 | 7.22E+06 | 5.32E+06 |
| **RABC1** | At1g43890 | protein transport [GO:0015031] | 0.01761352 | 1.15680619 | **1.75415385** | **0.21014717** | 1.81E+06 | 2.01E+06 | 2.00E+06 | 1.64E+06 | 2.07E+06 | 2.53E+06 | 1.34E+06 | 1.73E+06 | 2.32E+06 |
| **CTIMC** | At3g55440 | gluconeogenesis [GO:0006094]; gly | 0.01775177 | 0.86764506 | **1.75075831** | **-0.20482312** | 2.86E+07 | 2.53E+07 | 2.36E+07 | 2.48E+07 | 2.44E+07 | 2.67E+07 | 2.96E+07 | 2.82E+07 | 2.96E+07 |
| **At2g40290** | At2g40290 | | 0.01775316 | 0.79877542 | **1.75072425** | **-0.32413816** | 5.30E+06 | 5.26E+06 | 2.77E+06 | 3.97E+06 | 4.75E+06 | 4.85E+06 | 5.31E+06 | 5.98E+06 | 5.69E+06 |
| **MDAR5** | At1g63940 | response to cold [GO:0009409] | 0.01822283 | 1.05487859 | **1.73938413** | **0.07707696** | 3.03E+07 | 3.21E+07 | 2.22E+07 | 3.10E+07 | 3.37E+07 | 3.15E+07 | 2.97E+07 | 3.17E+07 | 2.98E+07 |
| **CRLK2** | At5g15730 | | 0.01835524 | 0 | **1.73623996** | **#¡NUM!** | 3.95E+06 | 2.64E+06 | 0 | 0 | 0 | 0 | 3.34E+06 | 2.73E+06 | 2.05E+06 |
| **SRK2G;SRK2H** | | | 0.01860114 | 1.06603234 | **1.73046035** | **0.0922512** | 1.55E+06 | 1.51E+06 | 1.23E+06 | 1.51E+06 | 1.55E+06 | 1.18E+06 | 1.42E+06 | 1.45E+06 | 1.11E+06 |
| **AT9943_LOCUS11431** | | | 0.01899596 | 0.83258781 | **1.72133883** | **-0.26432565** | 1.29E+06 | 1.31E+06 | 1.22E+06 | 1.14E+06 | 1.03E+06 | 1.05E+06 | 1.37E+06 | 1.19E+06 | 1.31E+06 |
| **HISN4** | At4g26900 | amino acid biosynthetic process [G | 0.01942498 | 1.05881386 | **1.71163938** | **0.08244898** | 1.60E+06 | 1.20E+06 | 1.14E+06 | 1.32E+06 | 1.33E+06 | 1.54E+06 | 1.26E+06 | 1.26E+06 | 1.44E+06 |
| **LIR1** | At3g26740 | circadian rhythm [GO:0007623] | 0.01997129 | 1.21123098 | **1.69959396** | **0.27647401** | 801291 | 811687 | 0 | 812739 | 799560 | 726370 | 699219 | 624753 | 606848 |
| **At5g05010** | At5g05010 | protein transport [GO:0015031]; re | 0.01999606 | 0.87787179 | **1.69905554** | **-0.18791783** | 6.32E+06 | 5.20E+06 | 5.67E+06 | 5.75E+06 | 5.06E+06 | 4.66E+06 | 6.56E+06 | 5.88E+06 | 5.17E+06 |
| **SLY1** | At2g17980 | protein transport [GO:0015031]; ve | 0.02003082 | 0.72659905 | **1.69830137** | **-0.46076861** | 620289 | 489058 | 757531 | 540770 | 483105 | 463365 | 704155 | 639343 | 703353 |
| **ALDH11A3** | At2g24270 | | 0.02003853 | 0.90913829 | **1.69813406** | **-0.13742834** | 1.65E+07 | 1.82E+07 | 1.68E+07 | 1.75E+07 | 1.77E+07 | 1.83E+07 | 1.96E+07 | 1.97E+07 | 1.96E+07 |
| **UGT73B2** | At4g34135 | flavonol biosynthetic process [GO:( | 0.02010833 | 0.17183532 | **1.6966241** | **-2.54090147** | 0 | 405543 | 0 | 225738 | 0 | 0 | 508869 | 461036 | 343783 |
| **AXX17_At3g25100** | | | 0.02130126 | 1.12615713 | **1.67159469** | **0.17140813** | 1.50E+06 | 1.41E+06 | 0 | 1.36E+06 | 1.40E+06 | 1.34E+06 | 1.21E+06 | 1.29E+06 | 1.15E+06 |
| **CPN60B2** | At3g13470 | protein refolding [GO:0042026] | 0.02153392 | 0.91061778 | **1.6668769** | **-0.13508246** | 7.08E+07 | 7.45E+07 | 6.09E+07 | 6.40E+07 | 7.33E+07 | 7.32E+07 | 7.27E+07 | 7.84E+07 | 8.01E+07 |
| **SBP2** | At4g14040 | polar nucleus fusion [GO:0010197] | 0.02195434 | 0.87817988 | **1.65847963** | **-0.18741162** | 2.50E+06 | 2.26E+06 | 2.80E+06 | 2.01E+06 | 2.18E+06 | 2.18E+06 | 2.39E+06 | 2.45E+06 | 2.41E+06 |
| **GSL-OH** | At2g25450 | glucosinolate biosynthetic process | 0.02204785 | 0.79555042 | **1.65663381** | **-0.32997472** | 2.85E+06 | 2.16E+06 | 1.52E+06 | 2.71E+06 | 1.72E+06 | 1.84E+06 | 3.39E+06 | 2.25E+06 | 2.24E+06 |
| **RPL22B** | | | 0.02227237 | 0.9700529 | **1.6522336** | **-0.04386467** | 2.25E+07 | 1.95E+07 | 2.85E+07 | 2.12E+07 | 1.91E+07 | 2.40E+07 | 2.19E+07 | 1.96E+07 | 2.49E+07 |
| **AXX17_At1g37100;C24_LOCUS3506;CGE2** | | | 0.02242383 | 1.31946357 | **1.6492903** | **0.39995152** | 1.20E+06 | 883165 | 902280 | 1.43E+06 | 1.44E+06 | 1.82E+06 | 1.05E+06 | 1.16E+06 | 1.34E+06 |
| **PGK3** | At1g79550 | glycolytic process [GO:0006096]; re | 0.0225677 | 0.94812291 | **1.64651273** | **-0.076854** | 8.18E+07 | 7.58E+07 | 5.28E+07 | 7.58E+07 | 7.64E+07 | 7.04E+07 | 7.97E+07 | 8.16E+07 | 7.35E+07 |
| **HMGB4** | At2g17560 | chromatin organization [GO:00063 | 0.02309165 | 0.7018268 | **1.63654503** | **-0.51081306** | 469048 | 408404 | 296129 | 239199 | 500083 | 280073 | 339732 | 673752 | 438947 |
| **PER50** | At4g37520 | hydrogen peroxide catabolic proce | 0.02343934 | 0.83481176 | **1.63005464** | **-0.26047718** | 818851 | 748270 | 447306 | 913441 | 674385 | 357065 | 1.01E+06 | 834820 | 489856 |
| **At5g44380** | At5g44380 | response to oxidative stress [GO:0( | 0.02461714 | 1.03452343 | **1.60876234** | **0.04896632** | 3.89E+06 | 5.39E+06 | 2.91E+06 | 4.72E+06 | 4.81E+06 | 3.91E+06 | 4.54E+06 | 4.65E+06 | 3.81E+06 |
| **PATL3** | At1g72160 | cell division [GO:0051301] | 0.02463664 | 0.96326035 | **1.60841856** | **-0.05400231** | 7.36E+06 | 7.47E+06 | 7.80E+06 | 7.56E+06 | 6.91E+06 | 7.07E+06 | 7.87E+06 | 7.24E+06 | 7.26E+06 |
| **At3g05070** | | | 0.02474712 | 0.8901949 | **1.60647539** | **-0.16780686** | 375959 | 430125 | 0 | 490877 | 461557 | 433895 | 529829 | 525102 | 502401 |
| **At3g49470** | At3g49470 | protein transport [GO:0015031] | 0.02476898 | 0.86016076 | **1.60609191** | **-0.21732177** | 2.51E+06 | 3.16E+06 | 2.21E+06 | 2.59E+06 | 3.03E+06 | 3.61E+06 | 3.20E+06 | 3.37E+06 | 4.15E+06 |
| **WDL5** | At4g32330 | cellular response to ethylene stimu | 0.02507173 | 1.36041574 | **1.60081575** | **0.4440476** | 348661 | 386343 | 532429 | 574395 | 462519 | 492547 | 482313 | 299654 | 342293 |
| **RPS1** | At5g30510 | | 0.02522082 | 0.85020506 | **1.59824074** | **-0.23411725** | 4.09E+06 | 3.98E+06 | 3.61E+06 | 3.46E+06 | 3.66E+06 | 3.81E+06 | 4.26E+06 | 4.33E+06 | 4.26E+06 |
| **At1g17220** | At1g17220 | | 0.0252231 | 1.21806705 | **1.59820159** | **0.28459356** | 2.56E+06 | 3.24E+06 | 2.34E+06 | 2.75E+06 | 2.58E+06 | 2.37E+06 | 2.19E+06 | 2.09E+06 | 2.05E+06 |
| **RPS8A** | At5g20290 | translation [GO:0006412] | 0.02569324 | 0.85487661 | **1.59018113** | **-0.22621189** | 2.83E+07 | 2.66E+07 | 2.65E+07 | 2.58E+07 | 2.49E+07 | 3.37E+07 | 3.01E+07 | 3.12E+07 | 3.74E+07 |
| **At5g38830** | At5g38830 | cysteinyl-tRNA aminoacylation [GO | 0.02596785 | 1.22755845 | **1.58556399** | **0.29579172** | 2.76E+06 | 2.03E+06 | 1.95E+06 | 2.11E+06 | 2.45E+06 | 2.92E+06 | 1.61E+06 | 1.88E+06 | 2.60E+06 |
| **TIM8** | At5g50810 | protein transport [GO:0015031] | 0.02632625 | 0.8304796 | **1.57961101** | **-0.26798336** | 227305 | 215755 | 330959 | 241034 | 278746 | 302144 | 283901 | 329831 | 375966 |
| **DHQS** | At5g66120 | amino acid biosynthetic process [G | 0.02718689 | 0.90345429 | **1.56564047** | **-0.14647648** | 1.64E+06 | 1.55E+06 | 1.54E+06 | 1.51E+06 | 1.49E+06 | 1.70E+06 | 1.73E+06 | 1.64E+06 | 1.83E+06 |
| **At1g03680** | At1g03680 | negative regulation of catalytic acti | 0.02748284 | 0.92161493 | **1.56093832** | **-0.11776401** | 2.20E+07 | 2.44E+07 | 1.84E+07 | 2.43E+07 | 2.35E+07 | 2.34E+07 | 2.59E+07 | 2.52E+07 | 2.61E+07 |
| **At3g47520** | At3g47520 | chloroplast organization [GO:00096 | 0.02817901 | 0.90706486 | **1.55007424** | **-0.14072237** | 4.16E+06 | 3.46E+06 | 4.47E+06 | 3.76E+06 | 3.54E+06 | 3.97E+06 | 4.07E+06 | 3.87E+06 | 4.49E+06 |
| **rpoB** | AtCg00190 | DNA-templated transcription elong | 0.02862216 | 0.8779084 | **1.54329763** | **-0.18785767** | 160239 | 161370 | 0 | 191053 | 193535 | 319973 | 216419 | 222376 | 363750 |
| **AT9943_LOCUS13297;At3g51000;F24M12.40** | | | 0.02881339 | 1.22285915 | **1.54040567** | **0.29025825** | 2.33E+06 | 2.73E+06 | 2.70E+06 | 3.20E+06 | 3.47E+06 | 3.38E+06 | 2.57E+06 | 2.68E+06 | 2.96E+06 |
| **PGDH1** | At4g34200 | embryo development ending in see | 0.02891025 | 0.92685926 | **1.53894814** | **-0.10957781** | 3.63E+07 | 3.21E+07 | 2.33E+07 | 3.28E+07 | 3.02E+07 | 2.87E+07 | 3.44E+07 | 3.26E+07 | 3.18E+07 |
| **FAB2** | At2g43710 | defense response [GO:0006952]; fa | 0.02900144 | 0.80728396 | **1.53758038** | **-0.30885186** | 1.21E+06 | 713519 | 591243 | 776732 | 977004 | 655205 | 914696 | 1.16E+06 | 908061 |
| **rpl22** | AtCg00810 | translation [GO:0006412] | 0.02905351 | 0.85009694 | **1.53680133** | **-0.23430074** | 2.47E+06 | 2.58E+06 | 3.39E+06 | 2.61E+06 | 2.85E+06 | 2.98E+06 | 2.94E+06 | 3.45E+06 | 3.54E+06 |
| **AN1_LOCUS23800;AT9943_LOCUS20590;AXX17_At5g33550;C24_L** | | | 0.0291416 | 0.94523045 | **1.53548665** | **-0.08126199** | 1.63E+06 | 1.32E+06 | 1.30E+06 | 1.40E+06 | 1.25E+06 | 1.32E+06 | 1.50E+06 | 1.32E+06 | 1.37E+06 |
| **C24_LOCUS24203** | | | 0.02965347 | 1.06549933 | **1.52792449** | **0.09152969** | 1.30E+06 | 1.60E+06 | 1.56E+06 | 1.48E+06 | 1.55E+06 | 1.40E+06 | 1.42E+06 | 1.43E+06 | 1.31E+06 |
| **SGPP** | At2g38740 | | 0.02972254 | 1.21916576 | **1.52691407** | **0.28589429** | 2.26E+06 | 1.66E+06 | 0 | 1.79E+06 | 1.48E+06 | 1.47E+06 | 1.57E+06 | 1.23E+06 | 1.08E+06 |
| **HSP70-8** | At2g32120 | response to heat [GO:0009408] | 0.02981141 | 1.38319108 | **1.52561745** | **0.46800047** | 518957 | 678891 | 574400 | 738633 | 677127 | 667323 | 480042 | 504701 | 521255 |
| **SYCO** | At2g31170 | cristae formation [GO:0042407]; cy | 0.0299094 | 0.75967501 | **1.52419237** | **-0.39654573** | 234838 | 300236 | 412041 | 248051 | 295626 | 258005 | 349207 | 393398 | 312691 |
| **IPMS1** | At1g18500 | L-leucine biosynthetic process [GO: | 0.0299912 | 0.83032126 | **1.52300612** | **-0.26825845** | 782476 | 649412 | 251358 | 471121 | 589907 | 510169 | 570710 | 668443 | 653123 |
| **FAP1** | At3g63170 | fatty acid metabolic process [GO:0( | 0.03025901 | 1.02133886 | **1.51914524** | **0.0304616** | 4.41E+06 | 5.62E+06 | 3.48E+06 | 5.19E+06 | 5.78E+06 | 4.42E+06 | 5.11E+06 | 5.63E+06 | 4.33E+06 |
| **HST** | At5g48930 | cell wall organization [GO:0071555] | 0.03100213 | 0.86193053 | **1.50860849** | **-0.2143565** | 3.02E+06 | 3.27E+06 | 2.03E+06 | 3.43E+06 | 2.61E+06 | 3.23E+06 | 3.95E+06 | 3.25E+06 | 3.56E+06 |

| | | | | | | | | | | | | | | | |
|---|---|---|---|---|---|---|---|---|---|---|---|---|---|---|---|
| **KAS2** | At1g74960 | cold acclimation [GO:0009631]; em | 0.03105772 | 1.14021328 | **1.50783044** | **0.18930371** | 2.08E+06 | 2.42E+06 | 1.47E+06 | 1.83E+06 | 1.64E+06 | 2.16E+06 | 1.52E+06 | 1.47E+06 | 1.95E+06 |
| **GAPC2** | At1g13440 | gluconeogenesis [GO:0006094]; gly | 0.03115202 | 0.72857332 | **1.50651375** | **-0.45685394** | 2.07E+08 | 1.78E+08 | 2.92E+08 | 1.63E+08 | 1.49E+08 | 2.17E+08 | 2.41E+08 | 2.27E+08 | 2.59E+08 |
| **CYP38** | At3g01480 | lateral root morphogenesis [GO:00 | 0.03128467 | 0.89724778 | **1.50466837** | **-0.15642164** | 3.37E+06 | 3.31E+06 | 2.03E+06 | 2.84E+06 | 3.09E+06 | 3.11E+06 | 3.30E+06 | 3.41E+06 | 3.36E+06 |
| **RD29A** | At5g52310 | circadian rhythm [GO:0007623]; hy | 0.03158777 | 1.20137586 | **1.50048105** | **0.26468759** | 656337 | 540165 | 0 | 796646 | 773683 | 717749 | 690677 | 599314 | 614557 |
| **At5g12240;MXC9.20** | | | 0.03164663 | 1.2847449 | **1.49967247** | **0.36148193** | 621481 | 562575 | 520919 | 751569 | 891378 | 490255 | 536667 | 765902 | 357840 |
| **G6PD5** | At3g27300 | glucose metabolic process [GO:000 | 0.03230868 | 0.95205118 | **1.4906808** | **-0.07088897** | 1.23E+06 | 765434 | 865169 | 1.03E+06 | 904072 | 1.06E+06 | 1.08E+06 | 972203 | 1.10E+06 |
| **CCT5** | At1g24510 | | 0.03263492 | 0.83848236 | **1.48631745** | **-0.25414766** | 4.39E+06 | 3.71E+06 | 2.74E+06 | 3.84E+06 | 3.40E+06 | 3.97E+06 | 4.45E+06 | 4.38E+06 | 4.52E+06 |
| **DDI1** | At3g13235 | proteolysis [GO:0006508] | 0.03328784 | 0.88573047 | **1.47771433** | **-0.17506034** | 8.63E+06 | 9.38E+06 | 4.68E+06 | 7.62E+06 | 8.54E+06 | 7.80E+06 | 8.95E+06 | 9.22E+06 | 8.87E+06 |
| **FBA8** | At3g52930 | gluconeogenesis [GO:0006094]; gly | 0.03403366 | 0.91202793 | **1.46809139** | **-0.13285009** | 3.63E+07 | 3.15E+07 | 2.65E+07 | 2.90E+07 | 3.01E+07 | 3.28E+07 | 3.29E+07 | 3.31E+07 | 3.47E+07 |
| **SDR4** | At3g29250 | | 0.03428293 | 0.89215696 | **1.46492206** | **-0.16463054** | 2.27E+06 | 1.86E+06 | 3.37E+06 | 1.95E+06 | 2.04E+06 | 2.17E+06 | 2.24E+06 | 2.34E+06 | 2.33E+06 |
| **CCT1** | At3g20050 | | 0.03474303 | 0.88785145 | **1.45913229** | **-0.17160979** | 3.36E+06 | 3.08E+06 | 4.05E+06 | 2.99E+06 | 3.03E+06 | 3.37E+06 | 3.50E+06 | 3.47E+06 | 3.62E+06 |
| **AGD2** | At1g60680 | | 0.03487884 | 0.73150707 | **1.45743792** | **-0.4510563** | 306953 | 1.05E+06 | 1.52E+06 | 989771 | 786284 | 865022 | 1.37E+06 | 1.17E+06 | 1.06E+06 |
| **RBP47A** | At1g49600 | cellular response to heat [GO:0034 | 0.03529947 | 0.20790788 | **1.45223182** | **-2.26598368** | 733379 | 993244 | 873087 | 424410 | 178874 | 0 | 926223 | 962226 | 1.01E+06 |
| **PKP2** | At5g52920 | fatty acid biosynthetic process [GO: | 0.03560799 | 0.81332233 | **1.44845257** | **-0.29810087** | 6.72E+06 | 4.94E+06 | 7.61E+06 | 5.23E+06 | 4.77E+06 | 4.23E+06 | 6.10E+06 | 5.66E+06 | 5.74E+06 |
| **RPS13** | At5g14320 | translation [GO:0006412] | 0.0357768 | 0.91852418 | **1.44639856** | **-0.12261039** | 2.54E+06 | 2.70E+06 | 3.59E+06 | 2.75E+06 | 2.92E+06 | 2.91E+06 | 3.09E+06 | 3.15E+06 | 3.09E+06 |
| **AN1_LOCUS2300;AT9943_LOCUS1882;ATEH2;AXX17_At1g22700;F** | | | 0.0358943 | 0.96985552 | **1.44497447** | **-0.04415825** | 1.47E+06 | 1.80E+06 | 1.30E+06 | 1.52E+06 | 1.61E+06 | 1.45E+06 | 1.56E+06 | 1.66E+06 | 1.51E+06 |
| **GAPB** | At1g42970 | glucose metabolic process [GO:000 | 0.03607381 | 0.89456932 | **1.44280797** | **-0.16073481** | 9.86E+07 | 8.28E+07 | 8.82E+07 | 8.47E+07 | 7.63E+07 | 9.45E+07 | 9.18E+07 | 9.01E+07 | 1.04E+08 |
| **EIF(ISO)4G1** | At5g57870 | | 0.03627047 | 0.87537252 | **1.44044685** | **-0.19203101** | 3.19E+06 | 3.06E+06 | 1.98E+06 | 2.89E+06 | 3.01E+06 | 2.60E+06 | 3.46E+06 | 3.33E+06 | 2.94E+06 |
| **SAL1** | At5g63980 | abscisic acid-activated signaling pat | 0.03639113 | 0.90405777 | **1.43900445** | **-0.14551313** | 6.08E+06 | 6.35E+06 | 5.14E+06 | 5.45E+06 | 5.25E+06 | 4.92E+06 | 6.15E+06 | 5.86E+06 | 5.26E+06 |
| **FREE1** | At1g20110 | abscisic acid-activated signaling pat | 0.03641753 | 1.08975733 | **1.43868948** | **0.12400691** | 2.96E+06 | 2.97E+06 | 1.75E+06 | 3.01E+06 | 3.17E+06 | 2.51E+06 | 2.75E+06 | 2.87E+06 | 2.37E+06 |
| **CAP1** | At4g34490 | actin cytoskeleton organization [GC | 0.03645516 | 0.8768535 | **1.43824097** | **-0.18959228** | 3.13E+06 | 2.85E+06 | 2.64E+06 | 2.53E+06 | 2.82E+06 | 1.78E+06 | 2.99E+06 | 3.08E+06 | 2.06E+06 |
| **AN1_LOCUS12233;F3L24.22;FES1A** | | | 0.03648798 | 1.08494745 | **1.43785018** | **0.11762517** | 3.20E+06 | 3.82E+06 | 3.24E+06 | 3.50E+06 | 3.83E+06 | 3.39E+06 | 3.26E+06 | 3.44E+06 | 3.18E+06 |
| **BAM5** | At4g15210 | response to herbivore [GO:008002 | 0.03676171 | 0.78417084 | **1.43460427** | **-0.35076009** | 905498 | 860092 | 817333 | 658777 | 950497 | 751964 | 934025 | 1.08E+06 | 993122 |
| **RPN8A** | At5g05780 | innate immune response [GO:0045 | 0.03791342 | 0.8436886 | **1.42120709** | **-0.24521748** | 3.95E+06 | 3.97E+06 | 4.36E+06 | 3.84E+06 | 3.70E+06 | 3.49E+06 | 4.30E+06 | 4.63E+06 | 4.15E+06 |
| **VHA-H** | At3g42050 | | 0.03860445 | 0.75805577 | **1.41336263** | **-0.3996241** | 1.85E+06 | 1.56E+06 | 2.66E+06 | 1.30E+06 | 1.43E+06 | 1.69E+06 | 1.89E+06 | 1.96E+06 | 1.98E+06 |
| **NMT1** | At5g57020 | embryonic shoot morphogenesis [G | 0.03868322 | 0.90451594 | **1.41247739** | **-0.14478217** | 2.06E+06 | 1.14E+06 | 1.97E+06 | 1.78E+06 | 1.43E+06 | 1.69E+06 | 1.92E+06 | 1.67E+06 | 1.82E+06 |
| **AN1_LOCUS1942;AXX17_At1g18960;At1g18070** | | | 0.03925806 | 0.87415211 | **1.40607118** | **-0.19404374** | 1.16E+07 | 1.17E+07 | 8.20E+06 | 1.01E+07 | 1.22E+07 | 9.32E+06 | 1.14E+07 | 1.33E+07 | 1.14E+07 |
| **BCCP2** | At5g15530 | fatty acid biosynthetic process [GO: | 0.04056187 | 1.10721681 | **1.391882** | **0.14693775** | 2.48E+06 | 2.33E+06 | 3.15E+06 | 2.77E+06 | 2.45E+06 | 2.88E+06 | 2.55E+06 | 2.08E+06 | 2.68E+06 |
| **CSY4** | At2g44350 | citrate metabolic process [GO:0006 | 0.04083778 | 0.89668963 | **1.38893785** | **-0.15731938** | 1.60E+07 | 1.54E+07 | 1.32E+07 | 1.32E+07 | 1.53E+07 | 1.36E+07 | 1.55E+07 | 1.65E+07 | 1.49E+07 |
| **At5g27470** | At5g27470 | cytoplasmic translation [GO:00021 | 0.04098349 | 0.8474786 | **1.38739105** | **-0.23875115** | 3.10E+06 | 3.02E+06 | 3.41E+06 | 2.73E+06 | 2.70E+06 | 3.15E+06 | 3.46E+06 | 3.14E+06 | 3.53E+06 |
| **TUBB4** | At5g44340 | microtubule-based process [GO:00 | 0.04130354 | 0.7919685 | **1.38401275** | **-0.33648505** | 3.60E+06 | 2.75E+06 | 3.10E+06 | 2.53E+06 | 1.50E+06 | 3.70E+06 | 3.09E+06 | 2.46E+06 | 4.21E+06 |
| **RPN11** | At5g23540 | protein catabolic process [GO:0030 | 0.04207604 | 0.82377385 | **1.37596517** | **-0.27967976** | 4.47E+06 | 3.36E+06 | 4.03E+06 | 3.26E+06 | 3.02E+06 | 3.92E+06 | 4.21E+06 | 3.82E+06 | 4.35E+06 |
| **CPX1** | At1g03475 | chlorophyll biosynthetic process [G | 0.04234577 | 1.0598407 | **1.37318992** | **0.08384744** | 1.73E+07 | 1.79E+07 | 1.51E+07 | 1.91E+07 | 1.92E+07 | 1.94E+07 | 1.77E+07 | 1.79E+07 | 1.88E+07 |
| **AN1_LOCUS22780;AXX17_At5g19740;KETCH1** | | | 0.0425221 | 0.74777719 | **1.37138533** | **-0.41931963** | 2.36E+06 | 2.17E+06 | 1.78E+06 | 1.85E+06 | 1.82E+06 | 1.83E+06 | 2.73E+06 | 2.28E+06 | 2.34E+06 |
| **PSAL** | At4g12800 | photosynthesis [GO:0015979] | 0.0428131 | 0.44173396 | **1.36842337** | **-1.17875035** | 458512 | 496122 | 1.12E+06 | 0 | 559146 | 316721 | 283227 | 856269 | 843297 |
| **BCA5** | At4g33580 | carbon utilization [GO:0015976] | 0.04293978 | 1.16212387 | **1.36714015** | **0.21676385** | 3.59E+06 | 3.59E+06 | 3.12E+06 | 4.55E+06 | 4.02E+06 | 4.74E+06 | 3.70E+06 | 3.64E+06 | 4.12E+06 |
| **MSBP2** | At3g48890 | | 0.04294253 | 0.77823968 | **1.36711239** | **-0.36171356** | 688603 | 774252 | 650352 | 609139 | 693900 | 453904 | 814513 | 789366 | 653707 |
| **UGD3** | At5g15490 | carbohydrate metabolic process [G | 0.04302792 | 0.85555123 | **1.36624965** | **-0.22507385** | 1.37E+06 | 897853 | 327026 | 929737 | 969486 | 777843 | 1.02E+06 | 1.14E+06 | 973764 |
| **rps5** | At2g33800 | adaxial/abaxial pattern specificatio | 0.04308466 | 0.74022281 | **1.36567734** | **-0.4339685** | 1.76E+06 | 1.21E+06 | 1.61E+06 | 1.39E+06 | 1.28E+06 | 915669 | 1.67E+06 | 1.67E+06 | 1.50E+06 |
| **TIF3F1** | At2g39990 | embryo development ending in see | 0.04328512 | 0.89879676 | **1.36366142** | **-0.15393318** | 2.73E+06 | 2.06E+06 | 1.17E+06 | 2.07E+06 | 2.16E+06 | 2.02E+06 | 2.24E+06 | 2.36E+06 | 2.35E+06 |
| **PSP** | At1g18640 | embryo development ending in see | 0.04367814 | 0.83972581 | **1.35973587** | **-0.25200976** | 2.66E+06 | 2.75E+06 | 1.03E+06 | 2.41E+06 | 2.24E+06 | 1.84E+06 | 2.97E+06 | 2.68E+06 | 2.09E+06 |
| **RBP45B** | At1g11650 | mRNA processing [GO:0006397]; re | 0.04405058 | 0.55936175 | **1.35604834** | **-0.83814649** | 1.90E+06 | 1.49E+06 | 1.85E+06 | 1.29E+06 | 1.26E+06 | 647261 | 2.28E+06 | 1.73E+06 | 1.70E+06 |
| **FOLB1** | At3g11750 | folic acid biosynthetic process [GO: | 0.04443566 | 1.16996154 | **1.35226835** | **0.22646111** | 945986 | 649743 | 0 | 809319 | 690367 | 406826 | 745505 | 608435 | 275611 |
| **TIF3E1** | At3g57290 | DNA-templated transcription initiat | 0.04501629 | 0.77727575 | **1.34663031** | **-0.36350159** | 1.25E+06 | 843469 | 474335 | 620364 | 669161 | 821280 | 899286 | 869553 | 946806 |
| **CNX1** | At5g20990 | auxin-activated signaling pathway [ | 0.04513929 | 0.90366821 | **1.34544532** | **-0.14613493** | 1.96E+06 | 1.81E+06 | 2.07E+06 | 1.85E+06 | 1.73E+06 | 1.68E+06 | 2.01E+06 | 2.00E+06 | 1.82E+06 |
| **BGAL10** | At5g63810 | carbohydrate metabolic process [G | 0.04524405 | 0 | **1.34443856** | **#¡NUM!** | 192000 | 257673 | 0 | 0 | 0 | 0 | 151867 | 86614.6 | 196645 |
| **CP31B** | At5g50250 | innate immune response [GO:0045 | 0.04529637 | 0.85564076 | **1.34393657** | **-0.22492289** | 6.52E+06 | 8.66E+06 | 8.08E+06 | 6.96E+06 | 7.86E+06 | 7.23E+06 | 8.02E+06 | 9.64E+06 | 8.11E+06 |
| **GLO2** | At3g14415 | oxidative photosynthetic carbon pa | 0.04529713 | 0.91251563 | **1.34392928** | **-0.13207882** | 1.26E+07 | 1.10E+07 | 1.45E+07 | 1.06E+07 | 1.08E+07 | 1.42E+07 | 1.17E+07 | 1.24E+07 | 1.50E+07 |
| | | | 0.04532578 | 0.77687324 | **1.34365475** | **-0.36424888** | 8.89E+06 | 9.44E+06 | 4.49E+06 | 9.18E+06 | 1.03E+07 | 7.52E+06 | 1.29E+07 | 1.21E+07 | 9.70E+06 |
| **At4g31480;At4g31490** | | | 0.04539642 | 0.80913193 | **1.34297836** | **-0.30555315** | 1.94E+06 | 1.19E+06 | 1.18E+06 | 1.58E+06 | 1.26E+06 | 1.23E+06 | 2.02E+06 | 1.58E+06 | 1.42E+06 |
| **MFP2** | At3g06860 | fatty acid beta-oxidation [GO:0006 | 0.04609964 | 0.94235831 | **1.33630246** | **-0.08565238** | 7.07E+06 | 6.91E+06 | 6.34E+06 | 7.06E+06 | 6.22E+06 | 6.40E+06 | 7.30E+06 | 6.76E+06 | 6.82E+06 |
| **At5g63440** | At5g63440 | circadian regulation of translation [ | 0.04617582 | 0.181956 | **1.3355854** | **-2.45833845** | 0 | 108577 | 135423 | 52029.5 | 0 | 0 | 117294 | 56339.5 | 112312 |
| **REC2** | At4g28080 | chloroplast localization [GO:001975 | 0.04674549 | 0.84863004 | **1.33026028** | **-0.23679235** | 1.71E+06 | 2.02E+06 | 283895 | 1.13E+06 | 1.31E+06 | 1.44E+06 | 1.46E+06 | 1.52E+06 | 1.59E+06 |
| **SRC2** | At1g09070 | cellular response to hypoxia [GO:00 | 0.04708628 | 0.75394975 | **1.32710565** | **-0.40745971** | 879461 | 1.25E+06 | 416897 | 851747 | 885956 | 270887 | 982537 | 1.11E+06 | 571743 |
| **CHLI1** | At4g18480 | chlorophyll biosynthetic process [G | 0.04715833 | 0.90019136 | **1.32644162** | **-0.15169637** | 8.22E+06 | 8.07E+06 | 6.43E+06 | 7.16E+06 | 8.38E+06 | 8.28E+06 | 8.37E+06 | 9.29E+06 | 8.81E+06 |
| **CYS6** | At3g12490 | defense response [GO:0006952]; h | 0.04737525 | 1.05398082 | **1.32444845** | **0.07584861** | 1.63E+07 | 1.85E+07 | 7.74E+06 | 1.75E+07 | 1.62E+07 | 1.61E+07 | 1.64E+07 | 1.51E+07 | 1.56E+07 |
| **PER71** | At5g64120 | defense response to fungus [GO:00 | 0.04751423 | 0.87582421 | **1.32317626** | **-0.19128677** | 2.98E+06 | 2.54E+06 | 1.78E+06 | 2.45E+06 | 2.62E+06 | 1.53E+06 | 2.86E+06 | 2.97E+06 | 1.71E+06 |
| **PTP1** | At1g71860 | defense response [GO:0006952]; in | 0.04756164 | 0.81971311 | **1.32274313** | **-0.28680902** | 1.06E+06 | 1.27E+06 | 1.06E+06 | 940023 | 1.16E+06 | 870604 | 1.11E+06 | 1.33E+06 | 1.19E+06 |
| **RPS3AB** | At4g34670 | translation [GO:0006412] | 0.0475927 | 0.84388596 | **1.32245963** | **-0.24488004** | 3.68E+06 | 2.87E+06 | 3.80E+06 | 2.63E+06 | 2.69E+06 | 4.42E+06 | 3.33E+06 | 3.02E+06 | 5.19E+06 |
| **AXX17_At3g01390;At3g02200;F14P3.15;F1C9.1** | | | 0.04782561 | 0.76469539 | **1.32033949** | **-0.38704292** | 591207 | 540045 | 0 | 416011 | 335591 | 392887 | 522355 | 503639 | 470666 |
| **GGAT1** | At1g23310 | glycine biosynthetic process [GO:00 | 0.04784538 | 0.88533822 | **1.32016002** | **-0.17569939** | 7.06E+06 | 6.97E+06 | 6.63E+06 | 6.78E+06 | 6.92E+06 | 7.53E+06 | 8.10E+06 | 7.75E+06 | 8.14E+06 |
| **RPS18A** | At1g22780; A | translational initiation [GO:000641 | 0.04803763 | 0.78712796 | **1.31841845** | **-0.3453299** | 2.77E+07 | 2.44E+07 | 3.41E+07 | 2.24E+07 | 2.16E+07 | 3.38E+07 | 3.09E+07 | 3.04E+07 | 3.76E+07 |
| **CPN10-2** | At2g44650 | protein folding [GO:0006457] | 0.04811429 | 1.08224273 | **1.31772592** | **0.11402411** | 2.18E+07 | 2.53E+07 | 2.48E+07 | 2.53E+07 | 2.63E+07 | 2.60E+07 | 2.29E+07 | 2.53E+07 | 2.36E+07 |
| **MS1** | At5g17920 | 'de novo' L-methionine biosyntheti | 0.0482479 | 0.88712537 | **1.31652159** | **-0.17279009** | 2.12E+08 | 1.98E+08 | 1.86E+08 | 2.04E+08 | 1.74E+08 | 2.20E+08 | 2.19E+08 | 2.08E+08 | 2.46E+08 |
| **VEP1** | At4g24220 | response to wounding [GO:000961 | 0.04841515 | 1.16596899 | **1.31501869** | **0.22152942** | 1.10E+07 | 9.36E+06 | 7.33E+06 | 1.21E+07 | 1.05E+07 | 1.02E+07 | 9.84E+06 | 9.24E+06 | 9.03E+06 |
| **PCS1** | At5g44070 | arsenite transport [GO:0015700]; d | 0.04915837 | 0.83412847 | **1.30840256** | **-0.26165849** | 2.77E+06 | 2.62E+06 | 2.11E+06 | 2.15E+06 | 2.14E+06 | 1.83E+06 | 2.74E+06 | 2.43E+06 | 2.16E+06 |
| **CCOAOMT1** | At4g34050 | coumarin biosynthetic process [GO | 0.04932389 | 0.87977213 | **1.30694267** | **-0.1847982** | 9.12E+06 | 6.93E+06 | 4.40E+06 | 6.83E+06 | 6.52E+06 | 7.81E+06 | 7.86E+06 | 7.84E+06 | 8.37E+06 |
| **SRP9** | At3g49100 | negative regulation of translational | 0.04965607 | 0.26014943 | **1.30402762** | **-1.94258754** | 462088 | 754028 | 1.03E+06 | 0 | 180597 | 670684 | 690597 | 1.35E+06 | 1.23E+06 |
