## Supplemental table S2-3 for "Chemical modulation of chloroplast de- and re-differentiation reveals a role for the SAL1-PAP retrograde pathway in facilitating plastid transitions"

### SUPPLEMENTAL TABLES

**Table S1. List of proteins showing differential abundance between X57 and X56 treatments.** See Excel file.

**Table S2. Transcripts per million (TPM) values for the indicated genes.**

| Gene | ID | Mock_1 | Mock_2 | Mock_3 | X57_1 | X57_2 | X57_3 |
|---|---|---|---|---|---|---|---|
| *IPT1* | AT1G68460 | 0.410 | 0.260 | 0.254 | 0.088 | 0.043 | 0.083 |
| *IPT2* | AT2G27760 | 9.011 | 9.581 | 9.394 | 7.900 | 8.077 | 7.225 |
| *IPT3* | AT3G63110 | 24.556 | 30.820 | 26.683 | 14.285 | 17.224 | 15.052 |
| *IPT4* | AT4G24650 | 0.000 | 0.000 | 0.000 | 0.000 | 0.000 | 0.000 |
| *IPT5* | AT5G19040 | 7.576 | 6.157 | 8.025 | 3.990 | 3.142 | 2.595 |
| *IPT6* | AT1G25410 | 0.000 | 0.000 | 0.000 | 0.000 | 0.000 | 0.000 |
| *IPT7* | AT3G23630 | 6.698 | 5.891 | 6.731 | 1.314 | 1.277 | 2.279 |
| *IPT8* | AT3G19160 | 0.000 | 0.000 | 0.000 | 0.000 | 0.000 | 0.000 |
| *IPT9* | AT5G20040 | 34.597 | 33.921 | 32.500 | 28.675 | 25.734 | 25.510 |
| *CYP735A1* | AT5G38450 | 2.248 | 2.339 | 2.747 | 1.249 | 0.666 | 0.641 |
| *CYP735A2* | AT1G67110 | 8.769 | 9.112 | 9.098 | 1.792 | 1.534 | 1.439 |
| *LOG1* | AT2G28305 | 2.088 | 2.913 | 1.078 | 1.583 | 1.835 | 1.721 |
| *LOG2* | AT2G35990 | 0.153 | 0.054 | 0.580 | 0.110 | 0.213 | 0.103 |
| *LOG3* | AT2G37210 | 0.547 | 0.836 | 0.439 | 1.046 | 0.698 | 0.734 |
| *LOG4* | AT3G53450 | 1.288 | 0.681 | 0.997 | 0.606 | 0.336 | 0.243 |
| *LOG5* | AT4G35190 | 0.134 | 0.237 | 0.278 | 1.109 | 0.468 | 0.181 |
| *LOG6* | AT5G03270 | 0.176 | 0.248 | 0.181 | 0.315 | 0.306 | 0.295 |
| *LOG7* | AT5G06300 | 9.973 | 8.431 | 12.313 | 15.640 | 12.691 | 11.389 |
| *LOG8* | AT5G11950 | 28.035 | 25.181 | 24.675 | 22.485 | 25.015 | 26.825 |
| *CKX1* | AT2G41510 | 1.333 | 1.192 | 1.469 | 0.892 | 0.836 | 0.746 |
| *CKX2* | AT2G19500 | 0.000 | 0.000 | 0.000 | 0.000 | 0.000 | 0.000 |
| *CKX3* | AT5G56970 | 0.619 | 0.393 | 0.852 | 0.355 | 0.474 | 0.956 |
| *CKX4* | AT4G29740 | 2.962 | 3.616 | 3.059 | 3.781 | 3.503 | 3.114 |
| *CKX5* | AT1G75450 | 0.866 | 1.000 | 0.840 | 1.468 | 1.179 | 1.084 |
| *CKX6* | AT3G63440 | 2.978 | 4.389 | 3.844 | 2.059 | 1.926 | 2.213 |
| *CKX7* | AT5G21482 | 29.970 | 29.196 | 27.425 | 14.800 | 13.189 | 15.350 |
| *UGT73C1* | AT2G36750 | 6.782 | 9.406 | 8.222 | 21.113 | 17.738 | 23.543 |
| *UGT85A1* | AT1G22400 | 29.050 | 34.797 | 35.123 | 7.943 | 11.465 | 12.467 |
| *UGT76C1* | AT5G05870 | 8.303 | 9.260 | 8.449 | 12.739 | 12.589 | 12.911 |
| *UGT76C2* | AT5G05860 | 10.054 | 7.258 | 7.656 | 14.250 | 12.153 | 9.910 |
| *GLK1* | AT2G20570 | 55.126 | 54.487 | 62.083 | 7.957 | 9.850 | 6.811 |
| *GLK2* | AT5G44190 | 128.886 | 121.610 | 110.570 | 76.062 | 77.644 | 65.067 |
| *GNC* | AT5G56860 | 30.049 | 30.452 | 33.168 | 20.306 | 17.859 | 17.451 |
| *CGA1* | AT4G26150 | 3.400 | 4.060 | 2.746 | 0.797 | 0.683 | 0.483 |
| *ANAC013* | AT1G32870 | 6.043 | 6.277 | 7.122 | 18.662 | 10.460 | 11.537 |
| *AOX1a* | AT3G22370 | 14.117 | 16.432 | 17.120 | 64.028 | 47.707 | 48.090 |
| *APX2* | AT3G09640 | 0.067 | 0.070 | 0.206 | 0.430 | 0.487 | 0.537 |
| *CPK32* | AT3G57530 | 16.658 | 22.189 | 21.807 | 26.306 | 23.688 | 25.871 |
| *RCD1* | AT1G32230 | 52.189 | 59.019 | 57.425 | 68.427 | 56.786 | 55.651 |
| *SOT12* | AT2G03760 | 7.224 | 5.445 | 8.051 | 29.569 | 17.812 | 19.783 |

**Table S3. Primer and G-Block sequences.**

| | RT-qPCR primers | |
|---|---|---|
| **Gene** | **Forward primer** | **Reverse primer** |
| *UBC21* | TCAAATGCACCGCTCTTATC | CACAGACTGAAGCGTCCAAG |
| *GLK1* | TTGGGTCTCCGATTCTCCCTAT | GCAACTGGCGGTGCTCTAAAT |
| *GLK2* | ATCATGGTCACTTCAGGCCTTTGC | ATTGTCTTGTGGGAACACCGATGC |
| *GNC* | TTTGCTCATGGCTCTGTCGT | TGACTCCTGATATTAGACCAACTGC |
| *CGA1* | GCGACAGCAGCAAACAACACACTA | GGCCTTCCTTTGCCTTATTCCACA |
| *LHCB1B1* | AGCAGAGGACTTGCTTTACC | CATAGCCAACCTTCCGTTCT |
| *CAB3* | TAGCGATGGAGGACTCGATTA | CCTTCAACAGCTCCCATCAA |
| *PETE1* | GTCACGGTCATACAATCCGATAA | CCCAACGCAGATACACCTAAA |

| | Gateway (cloning) primers | |
|---|---|---|
| **Gene** | **Forward primer** | **Reverse primer** |
| *GLK1* | GGGGACAAGTTTGTACAAAAAAGCAGGCTGGATG<br>TTAGCTCTGTCTCC | GGGGACCACTTTGTACAAGAAAGCTGGGTG<br>GGCACAAGACGCGGTCGGAG |
| *FBN7a* | GGGGACAAGTTTGTACAAAAAAGCAGGCTGGATG<br>GCATTGATCCAACATG | GGGGACCACTTTGTACAAGAAAGCTGGGTG<br>ACTGTTGTATTCAAGATTC |

| SAL1 G-Block |
|---|
| AAAAAA<u>CC**ATG**G</u>CTTATGAAAAGGAACTCGATGCAGCCAAAAAGGCTGCAAGCCTGGCAGCTCGTCTCTGTCAG<br>AAGGTGCAAAAAGCACTGTTGCAGAGCGATGTTCAGAGCAAATCAGATAAGAGCCCGGTGACGGTTGCAGATTA<br>TGGTAGCCAGGCTGTTGTGTCTCTCGTTCTGGAGAAGGAATTGAGTAGCGAACCGTTTTCCCTGGTAGCAGAGG<br>AAGATTCTGGAGATCTGAGAAAAGATGGAAGCCAGGATACCCTCGAGAGAATCACAAAACTTGTGAATGATACC<br>CTGGCAACAGAAGAGTCCTTCAACGGCTCTACCCTGTCCACTGATGACCTCCTGCGTGCCATTGATTGTGGTAC<br>CTCTGAAGGTGGTCCGAATGGGAGGCATTGGGTTCTCGATCCCATTGACGGTACGAAAGGTTTTCTCCGTGGAG<br>ATCAATATGCTGTGGCACTGGGTCTCCTGGAAGAAGGAAAAGTAGTTCTTGGTGTTTTGGCTTGTCCCAATCTT<br>CCGCTGGCGAGCATTGCCGGTAACAACAAGAATAAAAGCTCCTCTGATGAGATTGGTTGCTTGTTTTTTGCTAC<br>TATTGGTAGTGGTACCTACATGCAGCTTCTGGATAGTAAAAGCAGTCCGGTTAAAGTTCAGGTGAGCTCCGTTG<br>AGAATCCTGAAGAGGCAAGTTTTTTTGAATCATTTGAGGGCGCTCATTCATTACATGATCTATCTAGCTCGATT<br>GCAAACAAACTGGGCGTTAAAGCCCCGCCGGTCCGTATCGATTCTCAAGCAAAGTATGGAGCACTGAGCCGCGG<br>TGATGGTGCAATATATCTACGATTTCCTCACAAAGGTTATCGTGAAAAGATTTGGGACCATGTTGCAGGGGCAA<br>TTGTTGTTACCGAAGCTGGTGGCATCGTTACAGATGCCGCTGGTAAACCACTCGATTTCTCAAAAGGGAAATAC<br>CTCGATCTGGATACCGGTATCATTGTTGCAAATGAAAAACTGATGCCTCTACTGCTTAAAGCAGTTCGTGACTC<br>CATCGCTGAACAAGAAAAAGCATCTGCCCTG**TAA**<u>GAATTC</u>AAGCAA |
